# Systematic Profiling and Functional Characterization of Alternative Splicing in *C. elegans* Olfactory Learning

**DOI:** 10.64898/2026.06.10.731382

**Authors:** Maoting Chen, Min Wu, Bina Koterniak, Minghai Ge, Jingting Liang, John Calarco, Yun Zhang

## Abstract

Alternative splicing (AS) is prevalent in neuronal gene expression. However, its function in learning has not been systematically characterized. Here, by profiling pan-neuronal translatome changes during a *C. elegans* learning paradigm, we show that AS remodels expression of neuronal genes with critical function in learning at the genome-wide scale. Intriguingly, AS operates on a functionally distinct gene set from those showing transcript abundance changes, serving as a separate regulatory layer in response to experience. Specifically, a neuronally enriched worm ortholog of a mitochondrial DNA helicase *twnk-1* displays significant learning-associated AS changes, with both isoform types acting in a pair of sensory neurons to regulate learning with distinct functions. AS of *twnk-1* modulates a cell-nonautonomous signal from neuronal mitochondria to peripheral tissues to regulate physiological states critical for learning. These results establish AS as a systematic regulator of learning and mechanistically reveal how AS shapes physiological states to facilitate learning.

## INTRODUCTION

Learning enhances survival by reducing exposure to danger and increasing opportunities to find food and mates. Previous studies on mechanisms underlying learning have identified critical roles of experience-dependent changes in transcription^1–3^. However, co- and post-transcriptional regulation, which also significantly shapes gene expression to provide additional layers of learning regulation, remain less well understood.

Alternative splicing (AS) is a co/post-transcriptional process that generates multiple isoforms of mRNA from a single gene and, thus, increases the diversity of transcripts and proteins encoded by eukaryotic genomes^4^. Previous studies using high-throughput sequencing found that around 95% of human genes containing multiple exons are processed by AS to produce various mature mRNAs^5^, and the prevalence of AS in gene expression is also well demonstrated in other vertebrate and invertebrate animals^6^. In the nervous system, AS plays an important role in regulating development and function across species^7–9^ and AS of several genes have been shown to regulate learning^10–13^. Meanwhile, disruption to AS is implicated in the pathology of several neurological diseases that impair learning and memory^14–16^. These results together suggest a fundamental role of AS in learning and brain plasticity. Intriguingly, it was shown that changing neural activity alters AS patterns of various genes in different neuronal cell types^17,18^. These findings suggest a mechanism by which experience-dependent neural activities, a critical basis for learning, reshapes AS-mediated gene expression to regulate learning. However, the pattern of AS has never been systematically analyzed at the pan-neuronal and whole-genome scales with single-gene resolution in an *in vivo* learning paradigm. Therefore, our system-level understanding of how AS responds to experience to regulate gene expression and support learning remains preliminary.

Experience-dependent changes in gene expression regulate various cellular events and physiological processes important for learning, including modulation of synaptic strength and circuit connectivity^1,3^. Meanwhile, internal states also significantly impact an animals’ ability to learn. Various physiological states, including stress level^19^, internal endocrine state^20^ and nutritional state^21^, have been shown to enhance or inhibit learning. Interestingly, the nervous system can signal to peripheral systems to regulate physiological states, as demonstrated by the neural regulation of stress responses in mice^22^ and worms^23^, and the hypothalamic signals conferring hunger state in mice^24^. However, neuronal genes underlying brain-body communication have not been fully characterized and, more importantly, whether AS mediates the expression of these genes to alter physiological state that supports learning has not been addressed.

Here, we applied translating-ribosome affinity purification (TRAP) coupled with RNA-Seq^25,26^ to the entire nervous system of *C. elegans* to examine how the neuronal gene expression is remodeled at the levels of ribosome-associated transcript abundance (referred to as “transcription” hereafter) and alternative splicing during an aversive olfactory learning paradigm. Using this systematic analysis, we found that similar to transcription, alternative splicing is also globally engaged in the nervous system to regulate gene expression during learning. Interestingly, learning-associated changes in transcript abundance and alternative splicing pattern are not converging but instead are enriched for genes encoding distinct functions. In addition, using RNA sequencing as an unbiased and comprehensive method, we discovered the function of two types of splicing isoforms of *twnk-1*, a neuronally expressed ortholog of the mammalian mitochondrial DNA helicase, in regulating learning. We showed that the aversive learning process alters the levels of two types of *twnk-1* isoforms that are expressed in different neurons and play distinct roles in regulating whole-body physiological states to support learning. Together, our analysis provides a system-level characterization of AS in an *in vivo* learning paradigm and uncovers the function of a conserved mitochondrial DNA helicase in learning by regulating body-brain communication in a splicing-mediated and experience-dependent manner.

## RESULTS

### Aversive learning process globally regulates alternative splicing (AS) and transcript abundance across the nervous system

Previously, we showed that worms grown on the standard laboratory food *Escherichia coli* strain OP50 learn to reduce their naive preference for the odorants of pathogenic bacteria *Pseudomonas aeruginosa* PA14 after training on it for 4-6 hours^27^. Using this learning paradigm, we sought to characterize AS at the pan-neuronal and the whole-genome scales. We performed translating-ribosome affinity purification coupled with RNA-Sequencing (TRAP-RNAseq)^25,26^ to systematically compare the ribosome-associated mRNAs (‘translatome’) in the entire nervous system of the naive OP50-fed worms and the PA14-trained worms (Figure S1A and Methods). To exclude the AS events elicited by different food sources (*E. coli* versus *P. aeruginosa*), we included two mock-trained conditions where worms were trained on another two strains, PA14-*gacA(-)* and PAK, that generate reduced virulence^28^ and induce non-significant level of learning^27^. Compared with wild-type worms, the worm strain used for pan-neuronal TRAP-RNAseq experiment exhibited similar susceptibility towards (Figures S1B-D) and learning indices (Figures S1E and S1F) induced by the three *Pseudomonas* strains. Consistently, during TRAP-RNAseq experiment, we observed robust learning behavior in PA14-trained condition and non-significant behavioral changes in either PA14-*gacA(-)*- or PAK-trained condition for pan-neuronal TRAP worms (Figure S1G).

To analyze the translatome in naive, PA14-trained and mock-trained worms, we mapped all ribosome-associated transcripts, quantified ribosome-associated transcript abundance (referred as “transcription” hereafter), and identified and quantified all alternative splicing events defined as local splicing variations (LSVs, Methods) in each condition. In total, we identified transcripts of 18889 neuronally expressed genes and 6227 neuronal LSVs from 3645 genes detected in all conditions. Principal component analysis (PCA) plots using the abundance of all neuronally expressed transcripts or splicing levels of neuronal AS events showed clear separation of PA14-trained groups from the rest (Figures 1A and 1B), suggesting that PA14 exposure regulates gene expression globally at both transcript abundance and alternative splicing levels.

**Figure 1.**
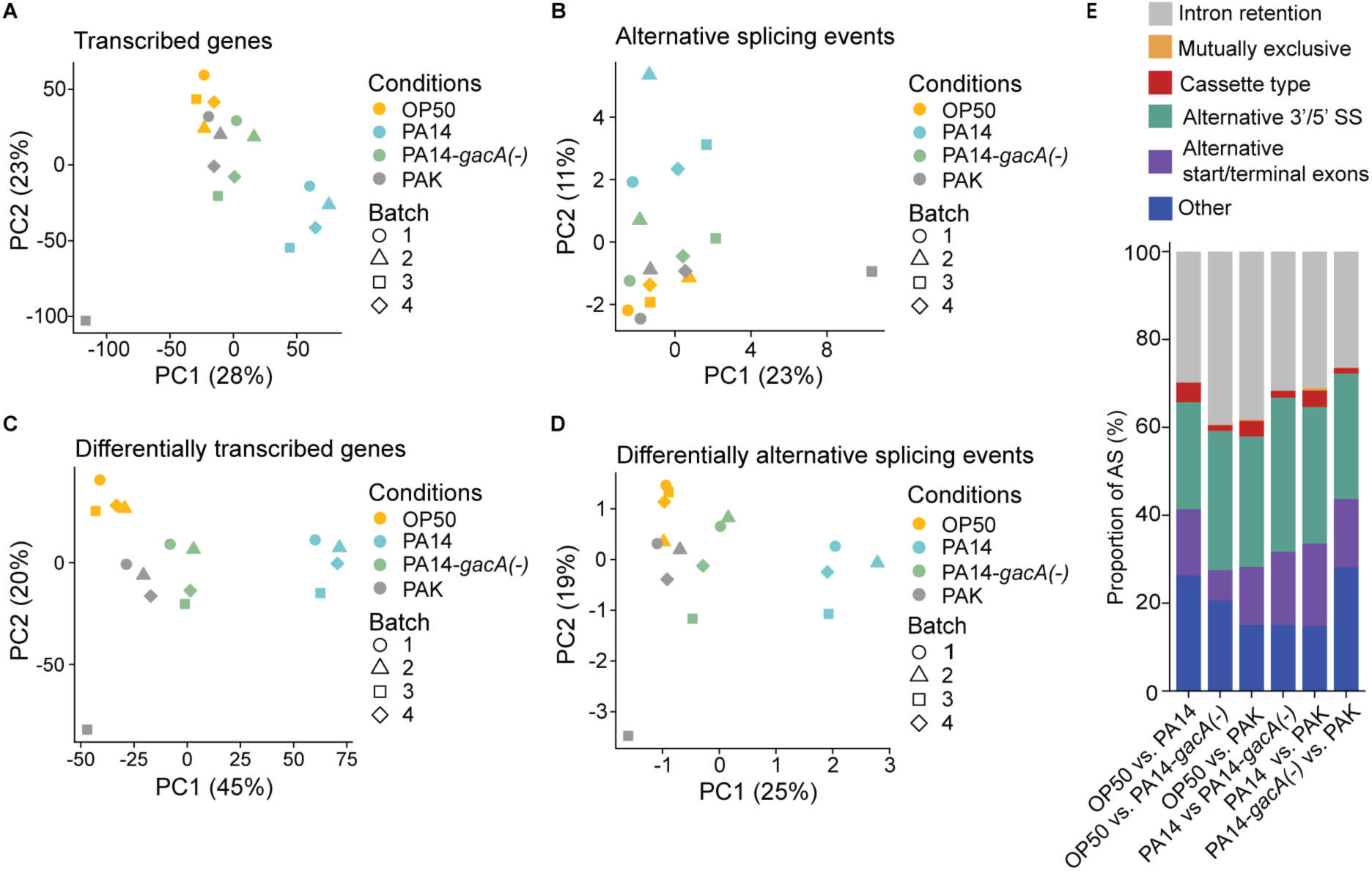
TRAP-RNAseq reveals both differential transcript abundance (DT) genes and differential alternative splicing (DAS) events upon PA14 training, also see Figure S1. (A and B) PCA analysis on all neuronally expressed genes and all neuronal AS LSVs shared by all samples, respectively. LSV, local splice variant, consists of splice junctions sharing the same donor or acceptor sites. (C and D) PCA analysis on DT genes (FDR < 0.05) and DAS LSVs (|dPSI| > 0.2, probability of change > 0.6) shared by all samples. dPSI, delta percent splice in, measures the difference of the frequency of one junction relative to all junctions within the same LSV between conditions. For A and C, PCA analysis used expression level of DT genes in each sample. For B and D, because one LSV contains multiple junctions, the PSI of the representative junction with the highest PSI variation across all samples for one LSV was used for PCA analysis. The expression level or PSI values were centered but not scaled across samples. (E) Relative proportion of DAS LSVs from six pair-wise comparisons classified into five major splicing types and an additional ambiguous type.

To characterize the types of splicing events generated under different training conditions, LSVs detected in any of the four conditions were additionally included for further analysis. The splicing types were classified by MAJIQ into five major categories^29^: intron retention, mutually exclusive splicing, cassette exon type, alternative 3ʹ/5ʹ splice site usage, and alternative start/terminal exons. Those that do not belong to the five types were defined as “Other”. We found that the neuronal AS events are distributed across the five types of splicing events, with 15%-30% of the events not categorized in any of the five types (Figure S2A). Consistent with previously identified splicing profile of the nervous system^29^, the major type of neuronal AS in *C. elegans* is alternative 3ʹ/5ʹ splice site usage.

We next evaluated potential batch effects of the sequencing samples by generating PCA plots with the potential outlier sample PAK3 removed (Figure S2B). Accordingly, we subsequently analyzed differential ribosome-associated transcript abundance (DT) and differential alternative splicing (DAS) in parallel with batch effect correction applied to DT analysis (Method). In total, we identified 7029 DT genes with FDR (false discovery rate) smaller than 0.05 (Table S1) and 767 DAS LSVs from 653 genes showing splicing change (change of PSI, percent splice in) larger than 0.2 from six pairwise comparisons (Methods) (Table S2). PCA plots using DT genes and DAS events showed clear clustering patterns of each training condition (Figures 1C and 1D). Again, the DAS events consist of the same five main splicing event types. Among these categories, the intron retention and alternative 3ʹ/5ʹ splice site usage are the predominant types of splicing events that are preferably regulated during exposure to different bacteria (Figure 1E). Together, the sequencing results demonstrate that both differential transcript abundance and differential AS are major regulatory mechanisms for gene expression in the PA14-induced olfactory learning paradigm.

### Motif analysis identifies candidate regulators of learning-associated DT genes and DAS events

We identified DT genes and DAS events that were significantly changed in PA14-trained group relative to the mock-trained groups as aversive learning-associated DT genes and DAS events (Methods). Using this analysis, we identified 1823 learning-associated DT genes (Table S3). For learning-associated DAS analysis, we specifically focused on the LSVs that were reliably detected in all four conditions and identified 160 learning-associated DAS events from 138 genes (Table S5).

Given the dynamic changes of learning-associated gene expression revealed by DT and DAS, we next sought to address how these changes were regulated. We first performed transcription factor (TF) binding motif analysis using XSTREME^30^ tool in MEME suite (Methods) on the learning-associated DT genes and identified enriched known and *de novo* motifs in up-regulated and down-regulated DT genes (Figure 2A). Meanwhile, we also quantified the changes in transcript levels induced by training with PA14 for all worm transcription factors (TFs)^31^ (Figure 2B). Using these two types of analyses, we found that the dominant class of the identified TF binding motifs contains a core GATA sequence, which is recognized by the GATA transcription factor family, including *egl-27*, several *elt* genes, and *end* genes^32,33^. Previous studies showed that *elt-2* regulates complex innate immune pathways to generate differential responses to different bacterial pathogens^32,34^, and *egl-27* mediates stress responses during aging^33^. Our analysis of DT genes also identified motifs recognized by ZTF-9, which was previously found to be involved in the aging process^35^ and a potential downstream target of the stress-responsive receptor DAF-12^36^. NHR-84 is another TF binding to one of the enriched motifs, and it is involved in wound repair^37^. Among all the matched TFs for the motifs we identified, DT analysis showed that *elt-2* and *elt-7* were significantly upregulated in the PA14-trained condition compared with the naïve condition while others were not (Figure 2B). These results together suggest a regulatory framework for aversive training-induced transcriptional changes.

**Figure 2.**
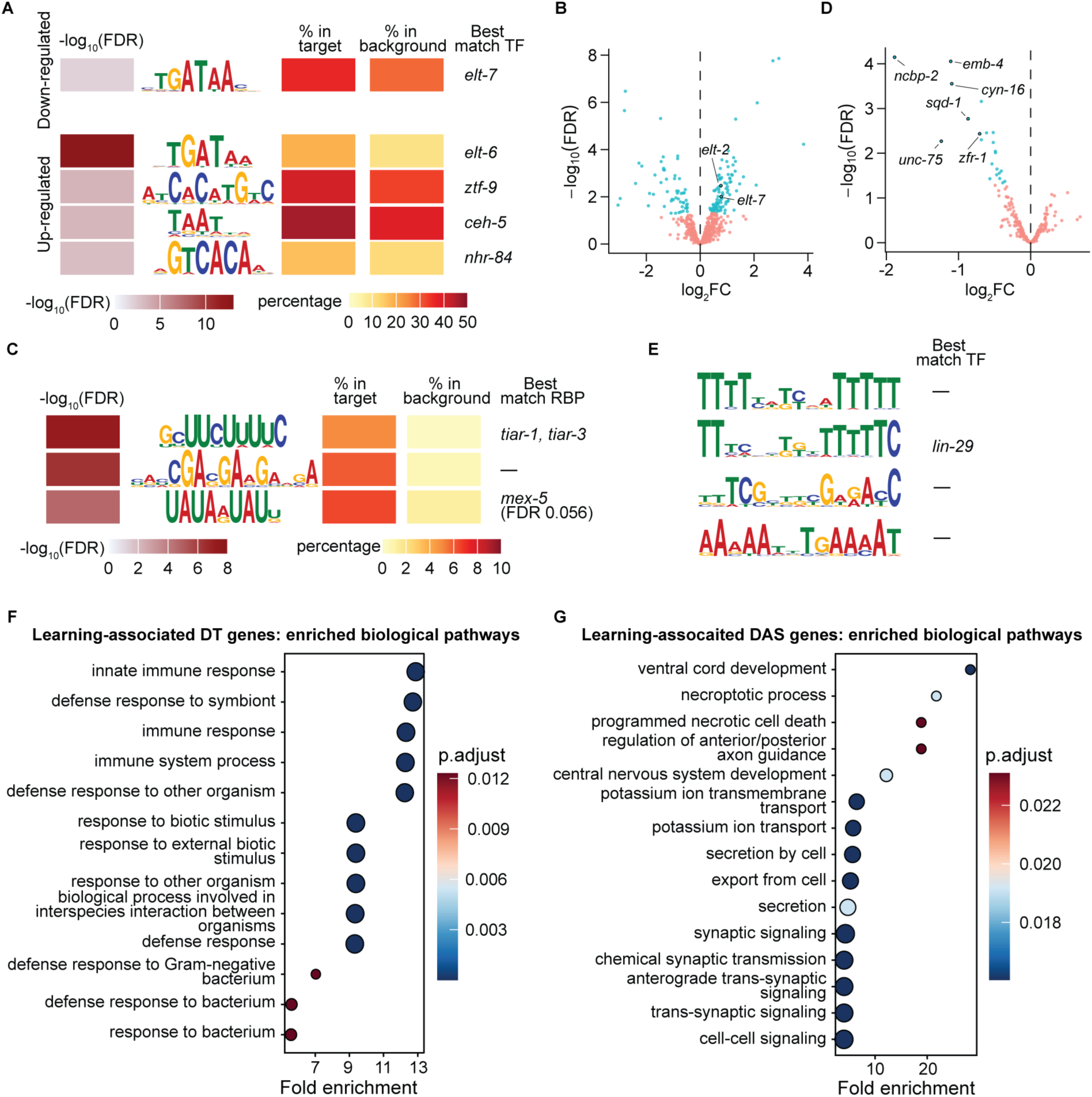
Differential AS and transcript abundance changes diverge in function during aversive learning with potential regulatory motifs identified, also see Figures S2 and S3. (A, C) Significantly enriched motifs identified in sequence regions around transcription start sites of learning-associated DT genes compared to those of neuronally expressed genes excluding learning-associated genes (A). Significantly enriched motifs identified in sequence regions around splice sites of learning-associated DAS events compared to those of neuronal AS events excluding learning-associated events (C). Motifs are presented as sequence logos with the height of the letters indicating relative frequency. The percentage of the motifs in the target and background groups, and the enrichment FDR (in −log_10_-transformed values) are displayed for each motif. If a motif matches with a known motif for a transcription factor (TF) or RNA binding protein (RBP) in *C. elegans*, the TF or RBP is shown. For similar motifs of the same group only the motif with the highest occurrence or strongest enrichment is shown (Methods). (B, D) Volcano plots showing gene expression difference of transcription factors (B) or splicing factors (D) between naive and PA14-trained conditions. Genes with FDR <0.05 are shown in blue, otherwise shown in pink. The transcription factors recognizing identified motifs with significant DT are labeled (B). The top 6 splicing factors with significant DT are labeled (D). (E) Motifs identified in promoter regions of downregulated splicing factors. Motifs are presented as sequence logos with the height of the letters indicating relative frequency. If a motif matches with a known motif for a transcription factor (TF) in *C. elegans*, the TF is shown. (F, G) Enriched biological pathways of learning-associated DT (F) and DAS genes (G) relative to all neuronal-expressed genes. Gene Ontology (GO) analysis was performed applying Benjamini-Hochberg (BH) method for multiple comparisons. The enrichment ratio is the frequency of input genes to the background frequency of genes in each GO term. Significantly enriched GO terms with p.adjust < 0.1 and/or top 15 enrichment ratios are shown.

We next characterized how DAS was orchestrated during learning by RNA-binding proteins (RBPs). We performed RNA-binding motif analysis on learning-associated DAS events (Methods). We identified enriched *de novo* motifs in DAS events and the conserved TIA-1/TIAR family of RNA-binding proteins TIAR-1 and TIAR-3 as potential RBPs recognizing one of the motifs (Figure 2C). The TIAR proteins are involved in the formation of processing bodies (PBs) and stress granules (SGs) in *C. elegans* for post-transcriptional mRNA regulations under stress conditions^38^. In addition, the homologs of TIAR-1 in flies and humans were found to regulate alternative splicing in processes such as apoptosis^39^. Together, these findings suggest a role of TIAR in pathogen-induced AS. We checked the PA14 training-induced transcript abundance changes of identified RBPs and found that *tiar-1* was significantly upregulated (Table S1). Consistent with the transcript abundance changes of identified TFs, not all potential TFs or RBPs regulating learning-associated DT or DAS genes showed altered transcript levels, suggesting that they could exert their regulatory role through mechanisms other than changes in their own transcript abundance.

We next examined differential transcript abundance of the splicing factors that are either directly or indirectly required for splicing after PA14 training. Interestingly, the splicing factors with significant transcript abundance changes are all downregulated, among which *unc-75* is neuronally enriched and known to regulate splicing in a context-dependent manner^40^ (Figure 2D and Methods). The downregulation of splicing factors might contribute to the relatively high fraction of intron retention observed in PA14 training-induced DAS events (Figure 1E). We further examined the motifs commonly presented in these downregulated splicing factors using MEME^41^ (Figure 2E). We identified several *de novo* motifs and a motif for the transcription factor LIN-29 that has been characterized for its role in cell fate determination via AS during development^42^. Together, our analyses reveal multiple regulatory layers potentially connecting aversive training experience to alternative splicing at the genome-wide scale.

### Differential AS and transcript abundance regulate experience-dependent gene expression through diverged functions and mechanisms

We next compared the functional effects of learning-associated DT genes and DAS events. To make the numbers of learning-associated DT genes and learning-associated DAS genes comparable and aid further analysis, we adjusted the thresholds for differential transcript abundance (Methods) and identified 144 learning-associated DT genes (Table S4). We observed that the learning-associated DT and DAS genes are expressed across different neuron classes including sensory neurons, interneurons and motor neurons based on CeNGEN single-cell RNA-seq expression dataset^43^ (Tables S4 and S5). We next performed Gene Ontology (GO) overrepresentation analysis^44^ using all neuronally expressed genes as the background. We found that these two layers of gene regulation preferentially regulate genes in different functions (Figures 2F, 2G, S2C and S2D). Learning-associated DT genes are only significantly enriched for the “non-motile cilium” in cellular components (Figure S2C) and are mostly enriched for biological pathways in immune and defense responses (Figure 2F), which is consistent with the identification of several stress-responsive TFs in the motif analysis (Figure 2A). In contrast, the learning-associated DAS genes are enriched in canonical biological pathways in the nervous system, such as synaptic signaling and axon guidance (Figure 2G), and enriched in cellular components of neurons, such as cation channel complex and somatodendritic compartments (Figure S2D). Given the prevalence of AS in regulating neuronal gene expression and function^7,29^, we further performed GO analysis on all neuronal AS genes against all neuronally expressed genes. Similar to learning-associated DAS genes, neuronal AS genes are also enriched in canonical neuronal biological pathways and cellular components (Figures S2E and S2F), suggesting that canonical neural functions are intrinsic for neuronal genes mediated by alternative splicing.

To further examine the divergence of DT and DAS, we characterized protein-protein interactions (PPI) of learning-associated DT and DAS genes using the STRING database^45^. We found more interactions among the learning-associated DT genes or the learning-associated DAS genes than between DT and DAS genes, although only a portion of genes have their interactions exhibited in the PPI network due to limited knowledge of PPIs in *C. elegans* (Figure S3A). Using a permutation test, we further confirmed the significantly enriched protein-protein interactions within the learning-associated DAS or DT gene groups, but not between the two gene groups (Figures S3B-S3D and Methods). Together, these results demonstrated that during the aversive learning process, genes showing differential AS and transcript abundance changes are enriched for different biological pathways and cellular functions.

### Experience-dependent AS regulates genes with critical functions in learning

Next, we sought to address how aversive experience-induced changes in AS regulate learning. We first selected the top learning-associated DAS events based primarily on three criteria: i. significant splicing change between naive and trained conditions but little change or a significant opposite change between naive and mock-trained conditions; ii. neuronal expression reported by the CeNGEN database for adult hermaphrodites^43^; iii. when applicable, homologs of mammalian genes whose functions in learning not full characterized. Applying these criteria, we focused on six learning-associated DAS events in six genes: the mutually exclusive exon event in *fbl-1*, which encodes a protein component in the extracellular matrix and regulates key developmental events such as organ morphology and cell migration^46,47^; the cassette exon event in *lips-5*, which is predicted to encode a lipase^48^; the alternative 3ʹ splice site event in *T24B8.7*, which is predicted to encode deubiquitinase activity^49^; the *de novo* alternative first exon event in *iff-1*, which encodes a translation initiation factor^50^; the alternative first exon event generating two types of isoforms (named as E1 and E6 isoforms) in *kcnl-2*, which encodes a calcium-activated potassium channel initially identified for its function in egg-laying^51^, and in *twnk-1*, which is predicted to encode a worm ortholog of a mitochondrial DNA helicase Twinkle in human^52^ (Figures 3A, 3B and S4).

**Figure 3.**
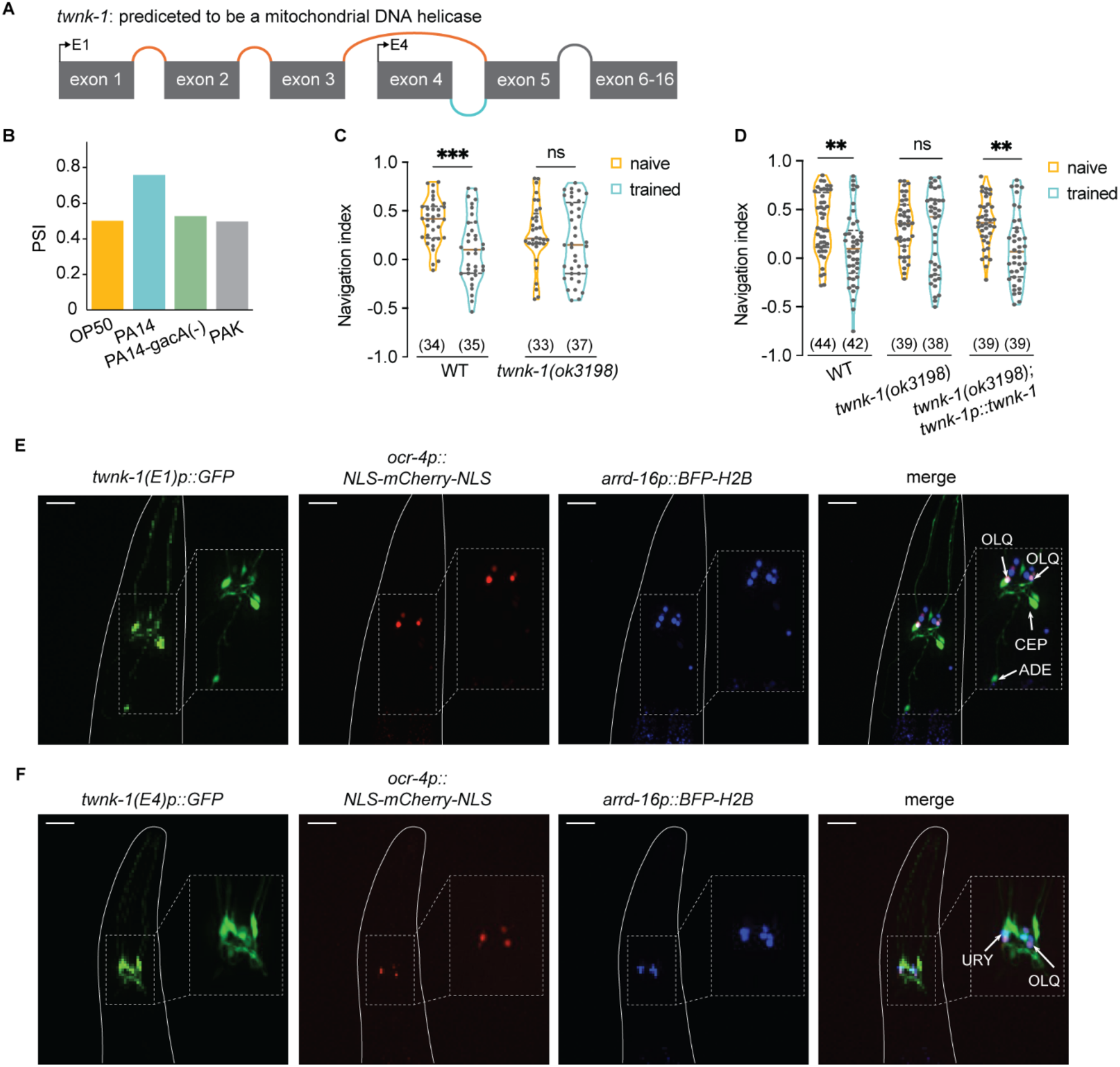
DAS gene *twnk-1* is expressed in the nervous system and regulates aversive olfactory learning, also see Figures S4, S5, S7A and S7B. (A) Schematic of the learning-associated AS event in *twnk-1*, with orange representing the splicing pattern downregulated, blue representing the splicing pattern upregulated after PA14 training and gray representing the shared splicing pattern. (B) PSI values of *twnk-1* splicing event in each condition. PSI, percent spliced in, measures the frequency of one junction relative to all junctions within the same LSV. (C) Violin plots of navigation indices towards PA14 supernatant in wild-type worms and *twnk-1(ok3198)* mutant worms under naive and PA14-trained conditions. (D) Violin plots of navigation indices towards PA14 supernatant in wild-type worms, *twnk-1(ok3198)* mutant worms, and transgenic *twnk-1(ok3198)* mutant worms carrying the full-length genomic *twnk-1* DNA under naive and PA14-trained conditions. (E and F) Sample images of co-localization of *ocr-4p::NLS-mCherry-NLS* and *arrd-16p::BFP-H2B* with *twnk-1(E1)p::GFP* or *twnk-1(E4)p::GFP* reporters in naive adult worms.Dashed lines outline the enlarged regions of interest. Lines outline the imaged worms. Arrows indicate the neurons with the neuron IDs specified. Scale bars, 20 µm. For (C and D), violin plots demonstrate individual data points and frequency distribution with horizontal lines illustrating median (brown solid) and quartiles (gray dashed). Two-way ANOVA across conditions within each genotype, with Sidak multiple comparison test. ** p<0.01, *** p < 0.001, ns, not significant. Numbers in parentheses indicate number of worms for each strain in each condition. Data was collected from 5 independent assays (C) and 6 independent assays (D).

To characterize the function of these genes, we first characterized their expression in the nervous system using GFP reporters. The transcriptional reporters driven by the 5ʹ regulatory sequences of these six genes all showed neuronal expression: *T24B8.7*, *kcnl-2*, and *iff-1* are broadly expressed across the nervous system, whereas *fbl-1*, *lips-5* and *twnk-1* are expressed in a limited number of neurons (Figures 3E, 3F, and S4). Next, we addressed the function of these genes by examining the learning phenotype in mutant harboring mutations in the genes.

We performed a crawling assay^53^ to test the deletion alleles in *T24B8.7*, *kncl-2* and *twnk-1*. We previously showed that naive adult worms are attracted to PA14 culture supernatant and navigate towards PA14 over a distance; training with PA14 decreases the navigation efficiency, indicating an experience-dependent reduction in olfactory preference^53^. We use a navigation index to quantify navigation efficiency towards PA14 supernatant throughout the assay, and higher preference for PA14 odorants generates a higher navigation index (Methods). We found that both *T24B8.7* and *twnk-1* mutants exhibited a learning-defective phenotype, as their navigation indices towards PA14 odorants showed no difference between naive and trained conditions, while wild-type control worms significantly decreased their navigation indices after training (Figures S4C and 3C). The *kcnl-2* mutant worms also did not exhibit a detectable training effect, although the navigation index of trained wild-type worms only showed a trend of decrease but was not significantly different from naive worms in this specific experiment (Figure S4D). We tested the *iff-1* mutant worms using a droplet assay^54^, where the preference between the odorants of OP50 and PA14 was measured in naive and PA14-trained swimming worms (Methods). We found that learning was defective in the *iff-1* homozygous mutant worms but remained intact in the heterozygous worms (Figure S4E). No behavioral characterization was performed on *fbl-1* or *lips-5*. Overall, our TRAP-RNAseq analysis uncovers the fundamental role of AS in globally regulating neuronal gene expression during learning and identifies genes with critical functions in learning that exhibit experience-dependent changes in alternative splicing.

### Aversive training alters splicing of *twnk-1,* a neuronally expressed mitochondrial DNA helicase

Given its role in regulating mitochondrial stress^52^ and its strong neuronal expression, we further investigated *twnk-1* splicing and functions. First, we found that expressing a full-length genomic DNA containing the regulatory and coding sequence of *twnk-1* rescued the learning defect in the deletion mutant worms, confirming the critical role of *twnk-1* in learning regulation (Figure 3D).

*twnk-1* encodes four annotated isoforms: an alternative first exon event generates two types of isoforms that differ in their DNA sequences at the 5ʹ end, and each of them consists of two isoforms distinct at the exon 12 region^49^. The learning-associated AS event detected for *twnk-1* is the alternative first exon event: one type of isoforms incudes exon 1-3 at the 5ʹ end and the other begins at exon 4 (Figure 3A) (respectively named as E1 and E4 isoforms thereafter). To determine the expression patterns of the E1 and E4 isoforms in the nervous system, we constructed GFP reporters using the corresponding promoters and identified the expressing neurons using neuron-specific red or blue fluorescent reporters. The E1 and E4 isoforms are expressed in partially distinct neuron sets: The E1 isoforms are mainly expressed in the dopaminergic neurons ADE and CEP as well as sensory neurons OLL and OLQ, while the E4 isoforms are expressed in the sensory neurons IL1, OLL, URY and OLQ and weakly in CEP (Figures 3E, 3F, and S5). The expression patterns we identified are consistent with the results reported by CeNGEN single-cell RNA-seq expression dataset^43^.

### *twnk-1* functions in OLQ sensory neurons regulates the neuronal activity and learning

Next, we further investigated where *twnk-1* acts in the nervous system to regulate aversive olfactory learning. To address this, we constructed neuron-specific rescue strains using a Cre/loxp-based inversion strategy^55^ (Figure 4A and Methods). In parallel, we examined the activity of *twnk-1* expressing neurons in response to the stimulation of PA14 supernatant in the context of OP50 supernatant using transgenic worms selectively expressing a genetically encoded calcium sensor, GCaMP6^56^, in these neurons (Methods).

**Figure 4.**
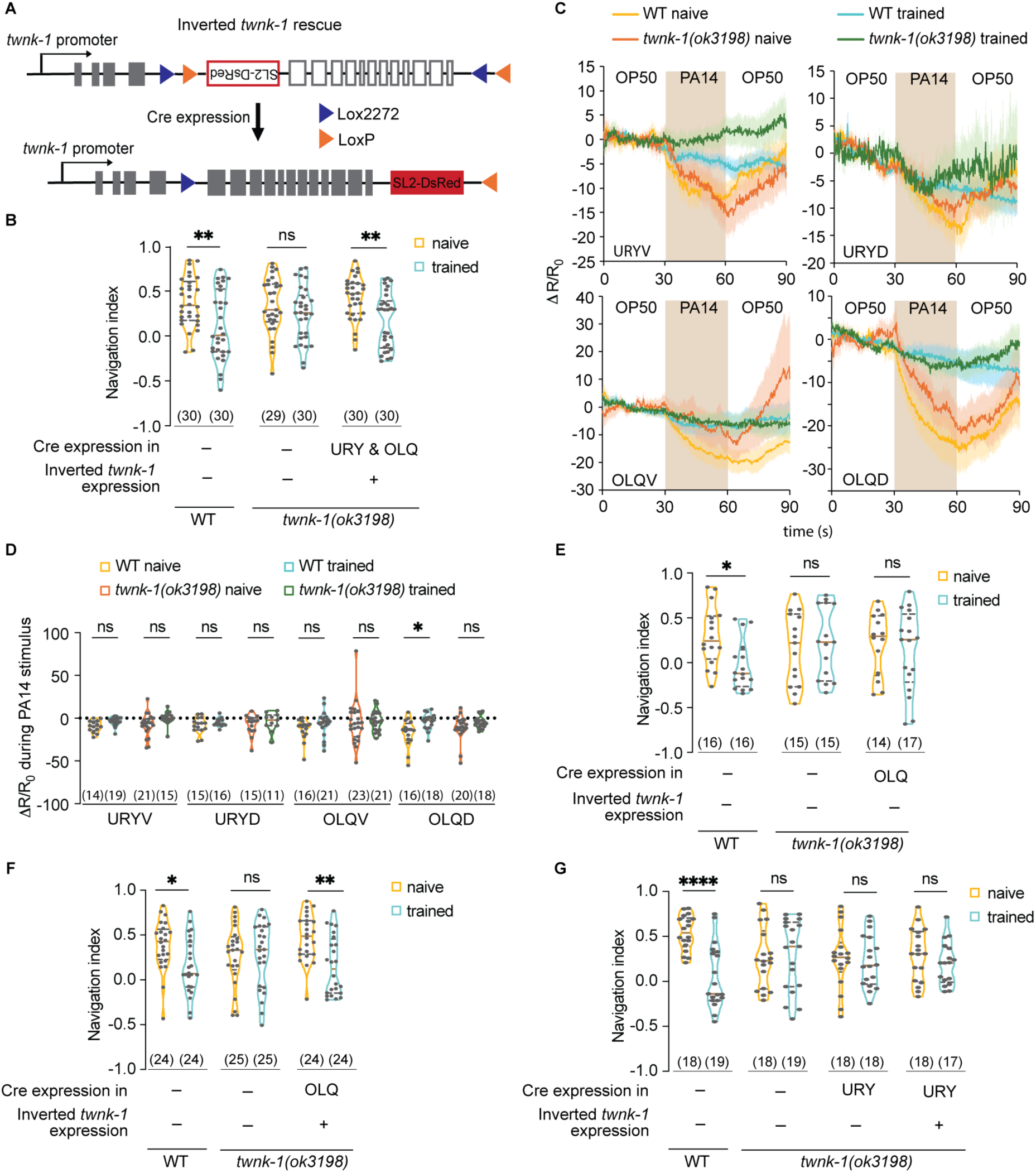
*twnk-1* acts in the OLQ neurons to regulate learning, also see Figure S6. (A) Schematic of neuron-specific *twnk-1* genomic rescue using Cre-loxp recombination. The gray filled boxes represent the *twnk-1* exons in the correct orientation. The gray unfilled box represents the *twnk-1* exons in the inverted orientation. The red filled and unfilled boxed represent the DsRed coding sequence and inverted DsRed coding sequence, respectively. The blue triangles represent the Lox2272 sites, and the orange triangles represent the LoxP sites. (B, E-G) Violin plots of navigation indices towards PA14 supernatant in naive and PA14-trained wild-type worms, *twnk-1(ok3198)* mutant worms, and transgenic *twnk-1(ok3198)* mutant worms with expression restored in different neurons. (C) Traces of GCaMP6 signals in URY and OLQ neurons in response to OP50-PA14-OP50 stimuli in wild-type and *twnk-1(ok3198)* mutant worms under naive and PA14-trained conditions. (D) Violin plots of quantification of average GCaMP6 signals across 30s PA14 stimulus in (C) for neurons in different genotypes under each condition. For (B, D-G), violin plots demonstrate individual data points and frequency distribution with horizontal lines illustrating median (brown solid) and quartiles (gray dashed). Two-way ANOVA across conditions within each genotype, with Sidak multiple comparison test (B, E-G); or multiple Mann-Whitney tests between naive and trained conditions across genotypes, with Holm-Sidak multiple comparison test (D). * p<0.05, ** p<0.01, **** p<0.0001, ns, not significant. Numbers in parentheses indicate number of worms assayed (B, E-G) or number of each neuron type recorded in each condition (D). Data were collected from 5 independent assays (B), 3 independent assays (D, E, G), and 4 independent assays (F). For (C), lines in traces, mean; shades in traces, s.e.m.; brown rectangular shade, PA14 stimulus. Λ′R/R_0_, see Methods.

First, we confirmed that no rescuing effect was detected in the control strains expressing only the inverted *twnk-1* cassette or only neuron-specific *cre* in the *twnk-1* mutant background, as the navigation indices in the naive and trained worms were comparable (Figures S6A, S6B, 4E and 4G). Next, we found that restoring *twnk-1* expression in the ADE and CEP dopaminergic neurons or in the OLL sensory neurons did not rescue the learning defect in the *twnk-1* deletion mutant animals (Figures S6C and S6D). Consistently, neither ADE nor CEP neurons showed significantly different responses to PA14 stimulation between naive and PA14-trained conditions in wild-type worms (Figures S6E and S6F). In addition, the OLL neurons exhibited PA14-evoked inhibition in the naive wild-type animals, which was eliminated after training. However, this training-dependent disinhibition remained intact in the *twnk-1* deletion mutant (Figures S6G and S6H), which is consistent with the non-rescuing effect on learning behavior in worms expressing OLL-selective *twnk-1*. The IL1 neurons did not show a calcium response to the PA14 stimulation either (Figures S6I and S6J).

Thus, we focused our analysis on the sensory neurons OLQ and URY. We found that expressing *twnk-1* in both OLQ and URY enabled significant training effects on navigation efficiency towards PA14 odorants and rescued the learning defect in the *twnk-1* deletion mutant animals (Figure 4B). At the level of neuronal activities, we found that among the URYV/D and OLQV/D classes, OLQD exhibited training-dependent changes in PA14-evoked calcium response, and this response was eliminated in the *twnk-1* mutant animals (Figures 4C and 4D). OLQV showed less robust and more variable responses to PA14 stimulation (Figures 4C and 4D). Prompted by the difference in the neuronal activities between OLQ and URY, we used an OLQ-selective promoter *ocr-4p* and a URY-selective promoter *tol-1p* to drive *cre* expression to restore *twnk-1* expression separately in these two classes of neurons. We found that the rescuing effect was only recapitulated in the OLQ-selective rescue strain but not in the URY-selective rescue strain (Figures 4F and 4G). Furthermore, the training-dependent effect on OLQD calcium responses to PA14 stimulation was restored in full-length genomic *twnk-1* rescued worms (Figures S6K and S6L). Together, these results indicate that OLQ neurons are the primary site where *twnk-1* acts to regulate learning, at least in part, by regulating OLQ neuronal activity.

### TWNK-1 isoforms regulate learning with distinct functions

We predicted the protein structures of E1 and E4 isoforms using AlphaFold4^57^ and found that the learning-associated DAS event resulted in a sequence of 40-50 amino acid difference at the N-terminal (Figures 5A and 5B). To investigate how the different N-terminal sequences influence functions of TWNK-1 isoforms during learning, we used CRISPR-generated *twnk-1(syb9885)* and *twnk-1(syb10064)* mutations that were respectively and specifically designed to knock out E4 and E1 isoforms (Figures 5C and 5D) and tested isoform-specific effects on learning using crawling assays. We found that both *twnk-1* E4 and E1 knockout mutants showed the learning defect that phenocopied the *twnk-1* deletion mutant worms (Figures 5E and 5F). These results suggest that the E1 and E4 isoforms are both required for the aversive learning of PA14.

**Figure 5.**
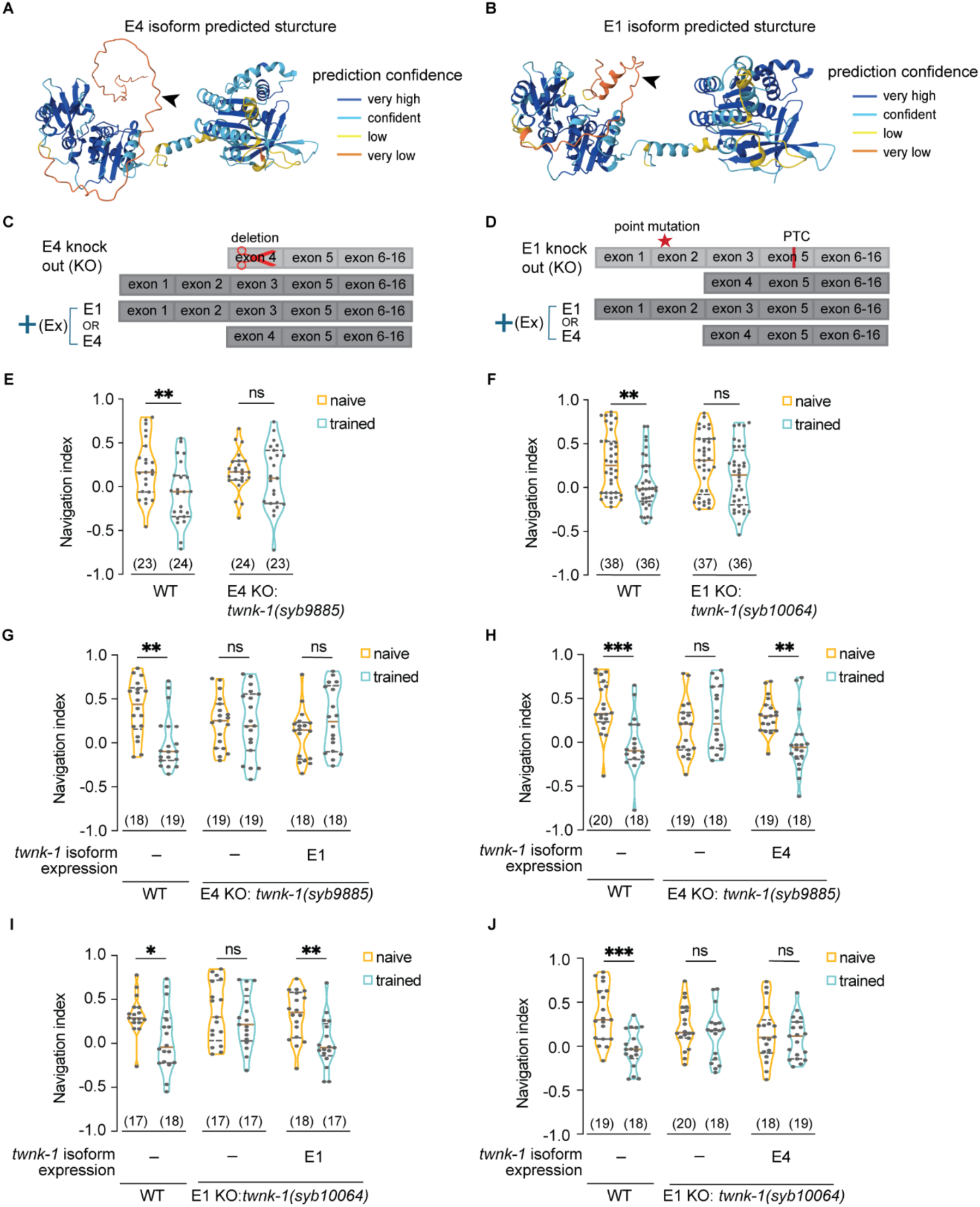
Both TWNK-1 isoforms are required for learning with distinct functions. (A, B) AlphaFold-predicted *twnk-1* E4 (A) and E1 (B) isoform (without exon 12) structures. Arrow heads indicate the regions where the learning-associated AS occurred. (C, D) Schematics showing the genetic background of the E4- or E1-specific knockout with E1 or E4 isoform-specific expression, respectively. Light gray indicates non-functional isoforms, and dark gray indicates functional isoforms. PTC, premature stop codon. Ex, extra-chromosome array. (E, F) Violin plots of navigation indices towards PA14 supernatant under naive and PA14-trained conditions in wild-type worms and *twnk-1(syb9885)* mutant worms (E); and in wild-type worms and *twnk-1(syb10064)* mutant worms (F). (G, H) Violin plots of navigation indices towards PA14 supernatant under naive and PA14-trained conditions in wild-type worms and *twnk-1(syb9885)* mutant worms (G, H), and transgenic *twnk-1(syb9885)* mutant worms expressing *twnk-1* E1 isoform cDNA expression driven by E1 promoter (G) or transgenic *twnk-1(syb9885)* mutant worms expressing *twnk-1* E4 isoform cDNA driven by E4 promoter (H). (I, J) Violin plots of navigation indices towards PA14 supernatant under naive and PA14-trained conditions in wild-type worms and *twnk-1(syb10064)* mutant worms (I, J), and transgenic *twnk-1(syb10064)* mutant worms expressing *twnk-1* E1 isoform cDNA expression driven by E1 promoter (I) or transgenic *twnk-1(syb10064)* mutant worms expressing *twnk-1* E4 isoform cDNA expression driven by E4 promoter (J). For (E-J), violin plots demonstrate individual data points and frequency distribution with horizontal lines illustrating median (brown solid) and quartiles (gray dashed). Two-way ANOVA across conditions within each genotype, with Sidak multiple comparison test. * p<0.05, ** p<0.01, *** p < 0.001, ns, not significant. Numbers in parentheses indicate number of worms for each strain in each condition. Data was collected from 4 independent assays (E), 6 independent assays (F), and 3 independent assays (G-J).

Two alternative hypotheses can explain the necessity of both E1 and E4 isoforms in learning: (1) both isoforms are necessary for learning with distinct functions; (2) alternatively, their functions may be similar, and it is the overall protein dosage that is critical for learning. To distinguish between these possibilities, we expressed E1 or E4 cDNA in the isoform-specific knockout mutants using the matched E1 or E4 promoter and performed crawling assays to evaluate if E1 isoforms could rescue the learning defect phenotype caused by E4-specific knockout, and vice versa (Figures 5C and 5D). We found that in the E4 knockout mutant worms, expressing E4 isoforms rescued the learning defect as expected while E1 isoforms did not rescue (Figures 5G and 5H). Conversely, in the E1 knockout mutant worms, expressing E1 isoforms rescued the learning defect phenotype as expected while E4 isoforms did not rescue (Figures 5I and 5J). Together with our findings showing the sufficiency of OLQ-specific expression of *twnk-1* genomic fragment in rescuing the learning defect in the *twnk-1* deletion mutant worms and the shared expression of E1 and E4 isoforms in the OLQ neurons, our results demonstrate that E1 and E4 isoforms are both needed with distinct functions in OLQ for the aversive learning.

### TWNK-1 AS coordinates mitochondrial stress and PA14-induced immune response to regulate aversive learning

The human and mouse orthologs of TWNK-1 are conserved mitochondrial DNA helicase that are essential for survival^58,59^. However, TWNK-1 in worms is dispensable^52^. Mitochondria provide the main energy source for cellular functions and have been shown to regulate neural plasticity^60,61^. Mitochondrial perturbation has also been implicated in many neurological diseases^61^. Thus, we first investigated the role of *twnk-1* in mitochondrial homeostasis. One common way to characterize mitochondrial stress is to measure the mitochondrial unfolded protein response (UPR^MT^), which is a transcriptional program activated upon stress^62^. The reporter *hsp-6p::GFP* is a target of this transcriptional program and widely used as a readout for mitochondrial stress^63^. We quantified the whole-body expression of *hsp-6p::GFP* in wild type, the *twnk-1* deletion mutant, and two isoform-specific knockout strains under naive and trained conditions. We found that the *hsp-6p::GFP* expression was significantly elevated under the naive condition in the *twnk-1(ok3198)* deletion and *twnk-1(syb9885)* E4-knockout animals, two independently generated mutant strains, indicating impaired mitochondrial homeostasis and induction of a mitochondrial stress response when TWNK-1 function encoded by the E4 isoforms is compromised (Figures 6A and 6B). In addition, the *hsp-6p::GFP* expression in PA14-trained worms was comparable across different genetic backgrounds (Figures 6A and 6B). These results together with the expression of *twnk-1* in the nervous system suggest that TWNK-1 regulates whole-body mitochondrial stress response in a cell-nonautonomous manner and reveal the different functions of E1 and E4 isoforms in this regulation. Increased UPR^MT^ has been shown to enhance resistance to pathogen infection in worms through mechanisms including regulation of innate immunity^23,64^. Therefore, we next asked whether *twnk-1* mutants exhibited higher resistance to PA14 by monitoring the survival of the worms on PA14 over three days (Methods). The *twnk-1* deletion mutant animals did show a significantly higher survival rate compared to wild type (Figure 6C). To further probe PA14-induced immune activation, we quantified expression of the immunity gene reporter *irg-1p::GFP* that was previously shown to increase in response to exposure to PA14^65^. Consistently, we found that training with PA14 increased *irg-1p::GFP* expression in all strains; however, the elevation was significantly reduced in *twnk-1* E4-specific knockout and deletion mutants but not in the E1-knockout mutant (Figures 6D and 6E), revealing a role of *twnk-1* E4 isoforms in regulating pathogen-induced immune activation.

**Figure 6.**
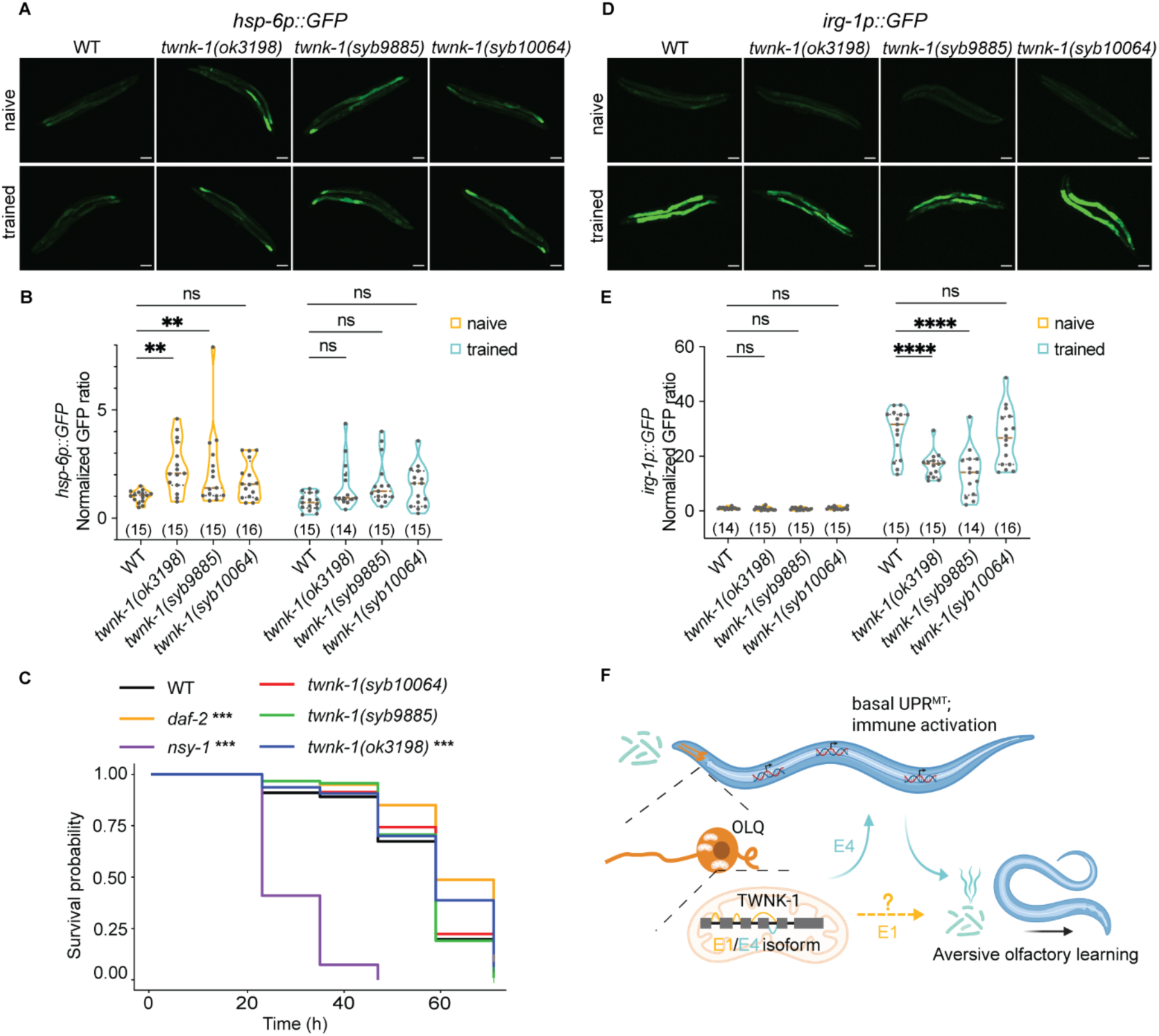
TWNK-1 AS coordinates mitochondrial stress and PA14-induced immunity. (A, D) Representative images of *hsp-6p*::GFP (A) and *irg-1p*::GFP (D) reporters under naive and PA14-trained conditions in the wild-type worms, *twnk-1(ok3198)*, *twnk-1(syb9885), and twnk-1(syb10064)* mutant strains. Images are maximum intensity projections, and brightness/contrast were applied equally using the same display parameters for each reporter. Scale bars, 100 µm. (B, E) Quantification of *hsp-6p*::GFP (B) and *irg-1p*::GFP (E) reporters under naive and PA14-trained conditions in the wild-type worms, *twnk-1(ok3198)*, *twnk-1(syb9885), and twnk-1(syb10064)* mutant strains. Violin plots demonstrate individual data points and frequency distribution with horizontal lines illustrating median (brown solid) and quartiles (gray dashed). Two-way ANOVA was used to compare mutant strains to wild type within each condition, with Dunnett multiple comparison test. * p<0.05, ** p<0.01, **** p<0.0001, ns, not significant. Numbers in parentheses indicate number of worms for each strain in each condition. Data was collected from 3 independent days. (C) Kaplan–Meier survival curve of wild-type, *twnk-1(ok3198)*, *twnk-1(syb9885), and twnk-1(syb10064)* mutant worms in a slow-killing assay of PA14 over three days. Pairwise Mantel–Cox log-rank tests compared each mutant strain with wild type, with experimental replicate included as a stratification factor and Bonferroni method as multiple comparison test. *** p < 0.001, ns, not significant. Summarized data from 3 independent assays are shown. 100 animals were used per genotype per assay (Methods). (F) Model of how *twnk-1* regulates aversive olfactory learning during PA14 exposure. Created in BioRender. Chen, M. (2026) https://BioRender.com/2rkzfk2.

Integrating the learning and mitochondrial stress phenotypes, we propose a working model for how AS of *twnk-1* contributes to learning (Figure 6F). *twnk-1* E4 isoforms are increased relatively to the E1 isoforms upon PA14 training. The E4 isoforms are required for full PA14-induced immune response, as loss of *twnk-1* E4 isoforms elevates UPR^MT^ in the naive condition, which in turn dampens PA14-induced immunity activation. This attenuation may weaken the experience-dependent association between PA14 odorants (conditioned stimulus) and PA14 infection-induced illness (unconditioned stimulus), thereby impairing aversive olfactory learning. In contrast, although the E1 isoforms are also required for learning, their absence does not affect PA14-induced immune response, suggesting that E1 may act through a distinct pathway—consistent with our conclusion based on the behavioral results that E1 and E4 isoforms are functionally distinct.

## DISCUSSION

Here, to our knowledge, we provide the first systematic view of how experience reshapes the neuronal translatome at two distinct regulatory layers by applying a genome-wide pan-neuronal ribosome-associated mRNA sequencing on naive and trained worms in an aversive learning paradigm. We further uncovered how a specific AS event regulates learning by characterizing the specific neuron where it is required, isoform usage dynamics, and physiological roles.

### Changes in alternative splicing and transcript abundance affect distinct sets of learning-associated genes

Our systematic analysis establishes alternative splicing as a critical mechanism underlying experience-dependent gene expression in learning by regulating isoform usage throughout the genome and across different neuronal cell types. The significant role of alternative splicing in regulating gene expression during learning is further underscored by the divergence of the gene pools primarily regulated by alternative splicing versus transcript abundance levels. Although transcript abundance changes also occur in genes contributing to neuronal functions, the most significantly regulated genes are enriched for defense responses after PA14 training; in comparison, genes exhibiting differential alternative splicing are primarily associated with canonical neuronal functions such as synaptic signaling, axon guidance, and transporter-associated events. Together with our analyses on protein-protein interaction and learning defects of the mutants carrying mutations in the DAS genes, these results support that AS mediates neuronal response to learning as a common gene regulatory process independent of transcript abundance regulation.

Similar regulatory divergence of transcription and AS is also found in other physiological processes in mammals. For example, it was shown that during the aging process, in the mouse hippocampus transcription regulated genes functionally enriched in the neuroinflammatory response while AS was associated with genes important for neural development and function, including synaptic transmission and plasticity^66^. Similarly, previous analysis of gene expression of different mouse tissues showed that transcription and AS contributed to tissue-specificity by regulating different sets of genes^67^. Together, these results reveal that it is a general mechanism that transcription and alternative splicing operate in distinct signaling pathways to generate different cellular responses to different internal and external conditions by regulating non-overlapping gene pools.

Intriguingly, our systematic analysis shows that while transcription factors are either increased or decreased by the aversive learning process, the expression of RBPs are ubiquitously decreased by training (Figures 2B and 2D). The decreased expression of RBP might be associated with the relatively large number of PA14 training-induced intron retention events. Increased intron retention has been shown to associate with Alzheimer’s disease and with the aging process in humans, mice and flies^68,69^, which suggests its importance in reshaping physiological functions and events. We propose that the mechanisms underlying different experience-dependent changes in transcription factors versus RBPs will provide insights into the divergence of these two types of regulators in controlling expression of genes with different functions.

### *twnk-1* reveals an isoform-specific mechanism for aversive learning

The learning-associated alternative first exon event in *twnk-1* uncovered two types of isoforms that are both necessary for learning with distinct physiological roles. The prediction on the protein structure suggest that alternative first exon in E1 and E4 isoforms leads to their distinct N-terminal structures that possess defining features of the intrinsically disordered region (IDR)^70^. IDRs have been widely found across species and represent polypeptides that are unable to generate any stably folded 3D structures^70^. IDRs have been shown to be subject to dynamic post-translational modification and alternative splicing, thus playing an important role in cellular signaling^70^, which provides plausible directions of how the *twnk-1* E1 and E4 isoforms differ from each other mechanistically.

Twinkle is the only identified mtDNA replicative helicase widely expressed in human and mouse mitochondria^58,59^. Inactivating Twinkle leads to mtDNA depletion and a series of mitochondrial diseases including neurodegenerative disorders^71,72^. In contrast, *C. elegans twnk-1* is expressed in a subset of neurons and its function as a mtDNA helicase is significant for base excision repair of mtDNA, ATP production and the larval-stage recovery from stressors that disrupt mtDNA^73^. Here, we further characterize the mitochondrial-specific functions of *twnk-1* based on UPR^MT^ induction in the *twnk-1* mutants, which is consistent with its colocalization with mitochondria^52,73^. Therefore, our results expand the function of *twnk-1* to the nervous system, identify the distinct functions of *twnk-1* isoforms in learning regulation, and, thus, connect *twnk-1-*mediated mitochondrial response to learning and memory (Figure 6F).

### Neuronal mitochondrial homeostasis modulates learning in a cell-nonautonomous manner

Mitochondria provide energy to support cellular functions and, thus, significantly impact many aspects of physiological states, such as metabolic states^74^, inflammatory responses^75^, and neuronal plasticity^76^. Malfunction of mitochondria is implicated in multiple neurological disorders^61^. Mitochondria also play a pivotal role in maintaining organismic functions under various stresses, many of which induce UPR^MT^ that restores cellular homeostasis^77^. However, sustained UPR^MT^ activation due to mutations or chronic stress could undermine cellular functions and disrupt normal physiological processes^78^.

In this study, we report that loss of *twnk-1* E4 isoforms modulates animals’ internal state by inducing peripheral intestine UPR^MT^ and suppressing infection-induced immune response, suggesting the cytoprotective and adaptive nature of *twnk-1*-induced UPR^MT^. These peripheral changes, in turn, potentially signal back to the neurons and regulate behavioral output. Periphery-brain communications have been implicated in neurological diseases where gut symptoms often manifest at the early stage of disease onset^79,80^. Overall, our findings support a new mitochondrial function of the neuronally expressed *twnk-1* in the brain-body-brain feedback. The systematic brain-body mitochondrial signaling enables efficient coordination of the whole organism in response to certain physiological changes^78^, and the feedback of body to brain facilitates the final behavioral modification.

Mechanistically, using the learning behavior assay we demonstrate that OLQ neurons are a key site of *twnk-1* function and that restoring *twnk-1* in OLQ is sufficient to rescue normal learning. Previously OLQ has been shown to regulate mechanical sensorimotor response by acting in an electrically connected mechanosensory circuit^81,82^. Here, our results suggest a new sensory modality of OLQ in olfaction. Although we have not directly tested whether loss of *twnk-1* in OLQ alone is sufficient to induce peripheral UPR^MT^ and alter immune responses towards pathogen exposure, our data raise the possibility that mitochondria dysfunction in a defined neuronal population can influence internal states and subsequent behavioral plasticity through cell-nonautonomous signaling. These findings propose a new component to the neuronal control of mitochondrial stress responses at the organismic level, in addition to the ASI-RIM TGF-β signaling pathway^83^, and provide insight into how neuronal metabolic state regulates associative learning by orchestrating systematic physiological responses in a cell-nonautonomous manner.

### Limitations of the study

First, our systematic AS profiling provides a list of potential learning-associated DAS genes. While our analysis that showes learning defect phenotypes in mutants carrying mutations in several of these genes supports important roles of DAS genes in learning, it is not practical to exhaust all the relevant genes and characterize the mechanisms underlying their functions in this study. Second, although we identified that the expression of *twnk-1* in OLQ is sufficient for aversive learning, we did not rule out the potential contribution of the function of *twnk-1* expressed in other neurons to learning. Finally, while our results on the function of *twnk-1* in aversive learning suggest a role of the human ortholog, Twinkle, in learning regulation, future studies in the mammalian system are needed to test this hypothesis directly.

## STAR * METHODS

### KEY RESOURCES TABLE

| REAGENT or RESOURCE | SOURCE | IDENTIFIER |
| --- | --- | --- |
| **Antibodies** |  |  |
| Mouse monoclonal anti-GFP | Memorial Sloan-Kettering Monoclonal Antibody Facility | cat# Htz-GFP-19F7, RRID:AB_2716736 |
| Mouse monoclonal anti-GFP | Memorial Sloan-Kettering Monoclonal Antibody Facility | cat# Htz-GFP-19C8, RRID:AB_2716737 |
| **Bacterial and virus strains** |  |  |
| *P. aeruginosa* PA14 | Tan *et al.*, 1999<sup>84</sup> | PA14 |
| *E. coli* OP50 | CGC | OP50 |
| **Chemicals, peptides, and recombinant proteins** |  |  |
| Protease Inhibitor Cocktail Tablets | Sigma-Aldrich | cat# 11836170001 |
| RiboLock RNase Inhibitor | Thermo Fisher Scientific | cat# EO0382 |
| NP40 | AG Scientific | cat# P1505 |
| Phosphate Buffered Saline, PBS (10X), pH 7.4 | Thermo Fisher Scientific | cat# AM9625 |
| Bovine Serum Albumin (IgG-Free, Protease-Free) | JacksonImmuno | cat# 001-000-162 |
| HEPES | Fisher Scientific | cat# AAJ16924AE |
| KCl (2M) | Thermo Fisher Scientific | cat# AM9640G |
| MgCl<sub>2</sub> (1M) | Thermo Fisher Scientific | cat# AM9530G |
| Cycloheximide | MilliporeSigma | cat# C7698 |
| DL-Dithiothreitol | MilliporeSigma | cat# D9779 |
| Recombinant Protein L | Thermo Fisher Scientific | cat# 29997 |
| Streptavidin magnetic beads | Thermo Fisher Scientific | cat# 65601 |
| TRIzol™ Reagent | Thermo Fisher Scientific | cat# 15596026 |
| Chloroform | MilliporeSigma | cat# 288306 |
| 2-Propanol | MilliporeSigma | cat# I9516 |
| Difco™ Agar Solidifying Agent | VWR | cat# 90000-762 |
| Bacto™ Peptone | Thermo Fisher Scientific | cat# 211677 |
| Sodium Chloride | MilliporeSigma | cat# S9888 |
| Potassium Phosphate Dibasic | MilliporeSigma | cat# P3786 |
| Potassium Phosphate Monobasic | MilliporeSigma | cat# P0662 |
| Calcium Chloride Dihydrate | MilliporeSigma | cat# 223506 |
| Magnesium Sulfate | MilliporeSigma | cat# M7506 |
| Cholesterol | VWR | cat# 97061-660 |
| Agar (for crawling assay) | MilliporeSigma | cat# 1.01614 |
| **Critical commercial assays** |  |  |
| OneStep RT-PCR Kit | QIAGEN | cat# 210212 |
| Illumina Stranded mRNA Prep kit | Illumina | cat# 20040532 |
| NEBuilder® HiFi DNA Assembly Master Mix | New England Biolabs | cat# E2621S |
| Q5® Site-Directed Mutagenesis Kit | New England Biolabs | cat# E0554S |
| E.Z.N.A.® Plasmid DNA Mini Kit I, (V-spin) | Omega Bio-tek | cat# D6943-02 |
| 2 × Phanta Max Master Mix (Dye Plus) | Vazyme | cat# P525-02 |
| Qubit™ RNA HS Assay Kit | Thermo Fisher Scientific | cat# Q32855 |
| **Deposited data** |  |  |
| pan-neuronal TRAP-RNAseq data | GEO | GSE333895 |
| **Experimental models: Organisms/strains** |  |  |
| *C. elegans*: Strain N2: wild isolate | Caenorhabditis Genetics Center (CGC) | N2 |
| *C. elegans*: strain JAC126: *csbls4* [*rgef-1p::eGFP* (+ intron)::*rpl-1* (spliced)::*unc-54* 3' UTR + *lin-15*] | Gracida *et al.*, 2017<sup>26</sup> | JAC126 |
| *C. elegans*: strain FX01052: T24B8.7 (*tm1052*) II | National BioResource Project | tm1052 |
| *C. elegans*: strain FX01245: T24B8.7 (*tm1245*) II | National BioResource Project | tm1245 |
| *C. elegans*: strain VC2138: *kcnl-2* (*ok2818*) I | CGC | VC2138 |
| *C. elegans*: strain JN215: *iff-1* (*tm483*) III/*hT2* [*bli-4*(*e937*) *let-?*(*q782*) *qIs48*] (I;III) | CGC | JN215 |
| Transgenic and mutant *C. elegans* strains | This paper | Table S6 |
| **Oligonucleotides** |  |  |
| Primers for cloning and PCR | This paper | Table S6 |
| **Recombinant DNA** |  |  |
| Plasmids for transgenic worms | This paper | Table S6 |
| **Software and algorithms** |  |  |
| MATLAB (R2024b) | MathWorks | https://www.mathworks.com/products/matlab.html |
| ImageJ (v1.54p) | National Institutes of Health | https://imagej.org |
| GraphPad Prism (v10) | GraphPad | www.graphpad.com |
| R (v4.5.1) | The R Project for Statistical Computing | https://cran.r-project.org/ |
| WormLab (v2024) | MBF Bioscience | https://www.mbfbioscience.com/products/wormlab/ |
| Cytoscape (v3.9.1) | Cytoscape Consortium | https://cytoscape.org/ |
| MEME Suite (v5.5.9) | The MEME Suite | https://meme-suite.org/meme/ |
| Custom code for DT and DAS analysis | This paper | github: https://github.com/Yun-Zhang-Lab/Pan-neuronal_TRAP-RNAseq.git |

### RESOURCE AVAILABILITY

#### Lead contact

Yun Zhang will respond to requests for information and resources.

#### Material availability

All strains and transgenes generated in this study are available from the lead contact upon reasonable requests without restriction.

#### Data and code availability

Pan-neuronal TRAP-RNAseq data have been deposited at NCBI Gene Expression Omnibus as GEO: GSE333895 and are publicly available upon the publication of the study.

All original code has been deposited at GitHub repository: https://github.com/Yun-Zhang-Lab/Pan-neuronal_TRAP-RNAseq.git.

### EXPERIMENTAL MODEL AND STUDY PARTICIPANT DETAILS

#### Animals and Maintenance

*C. elegans* hermaphrodite animals were maintained under standard conditions at 20-22 °C on nematode growth medium plates [NGM plates; 1.6% agar, NaCl (3 g/liter), Bacto Peptone (2.5 g/liter), 1 mM CaCl2, 1 mM MgSO4, 25 mM KPO4 (pH 6.0), and cholesterol (5 mg/liter)] containing a lawn of *Escherichia coli* OP50 cultured with Luria Broth overnight^85^.

#### Generation of transgenes and transgenic animals

All plasmids used were generated using Gateway LR recombination (Invitrogen #11791020) or Gibson assembly (New England Biolabs # E5520S).

To generate expression reporters for learning-associated DAS genes, the following 5ʹ regulatory regions were amplified from genomic DNA and cloned into an expression plasmid containing GFP converted from pPD95.77 (a gift from A. Fire): *fbl-1* (2k), *lips-5* (5.6k), *T24B8.7* (1.8k), *kcnl-2* (6k and 6.5k for E1 and E6 splicing isoforms, respectively), and *twnk-1* (1.4k and 3k for E1 and E4 splicing isoforms, respectively). For the *iff-1* expression reporter, the full-length *iff-1* genomic coding region (1.8k) was amplified from the genomic DNA, tagged with GFP, and then inserted in to the wild-type genome using miniMos technique^86^.

For cell identification and calcium imaging of *twnk-1*-expressing neurons, the neuron-selective reporters were generated by cloning the 5ʹ regulatory DNA regions of *arrd-16* (3k), *ocr-4* (4.8k), *flp-3* (1.5k), *cat-2* (1.5k) and *ser-2* (4.1k) into an expression plasmid containing a nucleus-localized BFP^87^ or mCherry^88^ or an expression plasmid containing GCaMP6^56^. The raw images of the co-localization between GFP reporters driven by *twnk-1* promoters and neuron-selective BFP or mCherry reporters are also shown (Figures S7A-S7H).

For *twnk-1* rescue experiment, the full-length *twnk-1* genomic DNA, including a 6.1k 5ʹ regulatory sequence, was amplified from the *C. elegans* genome and then cloned into pCR8 Gateway entry vector (Invitrogen K250020). The *twnk-1* E1 (1.7k) and E4 (1.8k) cDNAs were reverse-transcribed from *C. elegans* RNA and then cloned into pCR8 Gateway entry vector (Invitrogen K250020). *twnk-1* isoform-specific 5ʹ regulatory sequences and 3ʹUTRs were later inserted.

For neuron-specific rescue of *twnk-1* genomic sequence, the *cre* constructs were generated by cloning the neuron-selective 5ʹ regulatory sequences, *arrd-16* (3k), *ocr-4* (4.8k), *cat-2* (1.5k), *ser-2* (4.1k) and *tol-1* (5.5k), into the vector containing *cre* (a gift from C.I. Bargmann)^55^. A non-functional, inverted *twnk-1* construct was generated by Gibson Assembly on the inverted *dmsr-7* construct (a gift from C.I. Bargmann)^55^. Briefly, a genomic DNA fragment containing part of the *twnk-1* coding sequence and its 5ʹ regulatory sequence was first amplified and cloned at the 5ʹ side of the first Lox2272 and LoxP sites. The remaining *twnk-1* fragment was then inverted and inserted between the SL2-GFP and the second Lox2272 and inverted LoxP. At last, the GFP coding sequence was replaced by a DsRed coding sequence. The *twnk-1(ok3198)* mutant carrying the inverted *twnk-1* construct was crossed with the *twnk-1(ok3198)* mutant expressing the Cre recombinase under neuron-selective promoters. In the presence of Cre, the inverted *twnk-1* cassette together with DsRed is flipped into the correct orientation, enabling expression of functional TWNK-1 in the targeted neurons. The restoration of *twnk-1* expression was confirmed by DsRed expression in the expected neurons. For isoform-specific rescue of *twnk-1*, the 5ʹ regulatory region (1.4k) of the E1 isoforms was used for E1-specific expression. The 5ʹ regulatory region (3k) of the E4 isoforms containing CRISPR point mutations in the E1-specific knockout strain was used for E4-specific expression to avoid possible expression of E1 isoforms. The expression pattern of the mutated promotor of E4 isoforms was examined using a promoter driven GFP reporter and it was consistent with the wild-type promoter (Figure S7I). Since both types of E1 and E4 isoforms contains two sub-isoforms (differing by exon 12 inclusion), we expressed the mixture of the two variants for both types at the ratio based on TRAP-RNAseq data.

#### Aversive training and mock training

The aversive training with pathogenic bacteria PA14 and the mock training were performed as previously described^54^. The colony plates of OP50, PA14, and other bacteria, including the mock training bacteria with reduced virulence (PA14-*gacA(-)* and PAK), were prepared by streaking bacteria onto NGM agar plates, incubating at 37 °C overnight, storing at 4 °C and used within one week. Bacteria cultures were prepared by inoculating nematode growth medium [NGM; NaCl (3 g/liter), Bacto Peptone (2.5 g/liter), 1 mM CaCl2, 1 mM MgSO4, 25 mM KPO4 (pH 6.0), and cholesterol (5 mg/liter)] with single colonies from the bacteria plates and culturing at 26-27 °C overnight. OP50 or other bacterial culture (used within one week) was added to the 10-cm or 15-cm NGM training plates to cover the whole area, and extra liquid was removed. The plates were incubated at 27°C for 36-48 hours after the inoculating cultures were dried. On the assay day, young adult hermaphrodites were transferred to naive plates containing OP50, the aversive training plates containing PA14 or mock training plates containing PA14-*gacA(-)* or PAK after the plates were cooled to room temperature. After training for 4-6 hours at 20-22 °C, worms on naive plates, the training plates and the mock training plates were used for assays.

#### TRAP-RNAseq experiment

Translating ribosome affinity purification (TRAP) was performed as described^26^. Briefly, synchronized hermaphrodites that pan-neuronally express a GFP-tagged ribosomal protein RPL-1 were grown at 20–22 °C on NGM plates containing *E. coli* OP50 to young adult stage. Concentrated OP50 culture was added to the plates when needed to avoid starvation. Worms were washed off with M9 buffer [NaCl (5 g/liter), 1 mM MgSO4, KH2PO4 (3 g/liter), and Na2HPO4(6 g/liter)] and placed onto 15-cm NGM naive plates (*E. coli* OP50), training plates (pathogenic bacteria *P. aeruginosa* PA14) or mock training plates (*P. aeruginosa* strains with reduced virulence PA14-*gacA(−)* or PAK). After 4-6 hours of training, randomly picked worms were tested for learning in microdroplet assays and the remaining of worms were harvested to make lysates. Immunoprecipitation was performed to collect GFP-tagged ribosomes and associated RNA using anti-GFP antibodies (Rockefeller University catalogue no. Htz-GFP-19F7, RRID:AB_2716736; Htz-GFP-19C8, RRID:AB_2716737), Pierce™ Recombinant Protein L (Thermo Fisher Scientific catalogue No. 29997), and Invitrogen™ magnetic beads (Thermo Fisher Scientific catalogue No. 65601). cDNA libraries (4 conditions) were generated from the ribosome-associated RNAs using Illumina Stranded mRNA Prep kit (Illumina catalogue No. 20040532) and sequenced by Illumina NovaSeq.

#### Sequencing data analysis

Paired-end reads were mapped using STAR^89^ version 2.7.0e, *C. elegans* genome version WBcel235 and transcriptome annotation version WBcel235.106. Gene expression was quantified using HTSeq^90^ version 0.11.2. Differential gene transcript abundance was analyzed using edgeR^91^ version 4.8.2 with FDR < 0.05. Alternative splicing analysis was performed using MAJIQ^92^ version 3 with default parameters followed by additional stringent filters to obtain high-confidence AS and DAS events for further computational analysis. In MAJIQ, alternative splicing events are reported as local splice variants (LSVs), consisting of splice junctions sharing the same donor or acceptor sites. For each splice junction, splicing level is quantified as percent splice in (PSI), which measures the frequency of one junction relative to all junctions within the same LSV. PSI change (dPSI) and probability of change are quantified during differential AS analysis (|dPSI| > 0.2 and probability of changing > 0.6). We observed the separation of the samples across batches in the PCA plot with PAK3 removed for transcript abundance level but not in the PCA plot for splicing level (Figure S2B). Therefore, we performed differential gene transcript abundance with batch effect corrected.

To identify learning-associated DT genes or DAS events, we required the transcript abundance or splicing change to be significant between naive and PA14-trained conditions, but not between naive and mock-trained conditions; or significant between naive and PA14-trained and between naive and mock-trained conditions but with the difference between PA14-trained and mock-trained conditions also significant. For learning-associated AS analysis, LSVs that were reliably detected in all four conditions were focused. For learning-associated DT analysis, DT genes with FDR < 0.05 and |log_2_FC| > 1 from each pairwise comparisons were used in motif analysis. For GO and PPI analysis, to make the numbers of learning-associated DT and DAS genes comparable, a stringent threshold to DT genes were applied: FDR < 0.01, |log_2_FC| > 3.5. Except for MAJIQ analysis, all coding, statistical tests, and graphing were performed using R 4.5.1.

#### Motif analysis

For motif analysis on learning-associated DT genes (FDR < 0.05 and |log_2_FC| > 1) and DAS events, we used XSTREME^30^ in the MEME suite v5.5.9. To identify motifs responsible for the transcriptional regulation of learning-associated DT genes, the target sequences comprised the genomic region from 1000 nucleotides upstream to 100 nucleotides downstream of the transcription start site (TSS) for each candidate gene. The background sequences were the equivalent regions from all neuronally expressed genes, excluding the learning-associated DT genes. For motif analysis on learning-associated DAS events, the target sequences comprised the region from 100 nucleotides upstream to 50 nucleotides downstream of the donor or acceptor sites that are commonly shared by the junctions exhibiting learning-associated PSI changes within each LSV. The background sequences were the corresponding regions of donor or acceptor sites that are commonly shared by all junctions within background AS events, which include all neuronal AS events detected across all four conditions, excluding the learning-associated DAS LSVs. The sequences were extracted using R 4.5.1 and transcriptome annotation version WBcel235.106.

Within XSTREME, MEME^41^ and STREME^93^ were used for *de novo* motif discovery with motif window size from 6-15 bp. *C. elegans*-specific known motifs were queried from motif databases, Catalog of Inferred Sequence Binding Preferences (CISBP)^31^ and CISBP-RNA^94^ with matched transcription factor (TF) and RNA-binding protein (RBP) annotation, respectively. Simple enrichment analysis (SEA)^95^ was used to assess motif enrichment for known motifs and identified *de novo* motifs relative to background sequences. To annotate candidate regulators for each motif, TOMTOM^96^ was used to compare discovered *de novo* motifs with known motifs and identify the most matched known motif. To remove the redundancy of highly similar motifs, pairwise motif similarities were determined by TOMTOM^96^. Motifs with strong similarity (E-value <= 0.05) to the most occurring motif (seed motif) of a cluster were grouped together and clusters were merged if all motifs of one group were weakly similar (E-value <= 0.1) to the seed motif of another group. Default parameters were selected.

Below filters were applied to identify regulatory motifs for learning-associated DT genes and DAS events: 1. Motifs are significantly enriched (false discovery rates q-values <= 0.05) in the target sequences; 2. *de novo* motifs discovered by MEME or STREME are significantly occurring (E-values <= 0.05); 3. Non-significantly occurring STREME motifs are included if they show robust similarities to known motifs via TOMTOM (false discovery rates q-values <= 0.05); 4. To remove the similar motifs in each motif cluster, a representative motif with the most significant occurrence (lowest E-value) was selected.

We defined splicing factors based on Gene Ontology annotation^97^. Specifically, genes annotated to the GO terms mRNA splicing, via spliceosome (GO:0000398), spliceosomal complex (GO:0005681), or RNA splicing (GO:0008380) were classified as splicing-related genes and included in the analysis. For motif analysis on down-regulated splicing factors, MEME was used to identify the motifs within sequences spanning 1,000 nucleotides upstream to 100 nucleotides downstream of the TSS for each gene.

#### Gene ontology (GO) and protein-protein interaction analysis

We used R package clusterProfiler version 4.18.4^44^ and *C. elegans* annotation database version 2.1^97^ to perform GO overrepresentation analysis for enriched biological pathways and cellular components on learning-associated DT (FDR < 0.01 and |log_2_FC| > 3.5) and DAS genes. Sample datasets were learning-associated DT or DAS gene lists we identified or neuronal AS genes common in all four conditions, and the background gene set is all neuronal-expressed genes identified by edgeR. Bonferroni-Hochberg multiple testing correction was applied.

We used STRING App version 2.2.0^98^ in Cytoscape version 3.10.4^99^ to obtain the protein-protein interaction network of learning-associated DT and DAS genes. Permutation test was performed to confirm the interaction enrichment, where gene names of each node in the network were shuffled 1e+5 times and relative proportion distributions of DAS-DAS, DT-DT, and DAS-DT interactions across the shuffled networks were plotted. And the p-values for these DAS-DAS, DT-DT and DAS-DT interactions were calculated. The significance threshold 0.05 was used. All coding, statistical tests, and graphing were performed using R 4.5.1.

#### Microdroplet assay

The microdroplet assay was performed essentially as described^54^. Naive and trained worms were washed in nematode growth buffer [NGM buffer; NaCl (3 g/liter), 1 mM CaCl2, 1 mM MgSO4, and 25 mM KPO4 (pH 6.0)] briefly and then individually placed in twelve 1.3µL microdroplets of NGM buffer arranged in 3 (row) x 4 (column) layout on the surface of a sapphire window. Each microdroplet contains a single freely-swimming worm. Fresh cultures of OP50 and PA14 or mock training bacteria were prepared by inoculating single bacteria colonies in NGM medium and incubating at 26 °C with shaking at 125 rpm overnight. The sapphire window was then placed in an enclosed chamber and worms were exposed to 12 cycles of 30s air stream odorized with 20mL of OP50 cultures (OD = 0.3 – 0.4) alternating with 30s air stream odorized with 20mL culture of training bacteria (OD = 0.6 – 0.75). The flow of the air streams was controlled by solenoid valves and LabVIEW (National Instruments, Austin, TX). The movements of worms were recorded with a camera (Watex) and later analyzed using an algorithm in MATLAB (MathWorks, Natick, MA) that detects omega turns exhibited by each animal based on the body shape^100^. The olfactory preference of each worm between OP50 versus training bacteria was represented by the choice index calculated from the frequency of omega turns during alternating odorant exposure. As defined below, the value of choice index value is positively correlated with a worm’s olfactory preference for training bacteria and the learning index is the difference of choice indices between naive and trained worms.

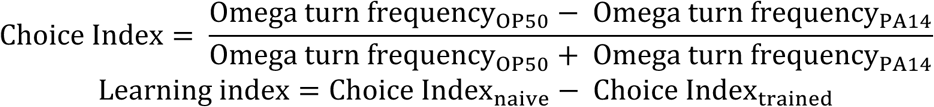

#### Single-worm crawling assay

Single-worm crawling assay towards PA14 odorants was performed as described^53^. 10-cm NGM assay plates [NaCl (3 g/liter), 1 mM CaCl2, 1 mM MgSO4, 25 mM KPO4 (pH 6.0), 1.6% agar, and cholesterol (5 mg/liter)] were prepared using 8 mL medium per plate 1 day before the assay. Fresh culture of PA14 (OD = 0.6 – 0.75) was centrifuged at 3,000rpm for 14 min and the PA14 supernatant was collected and diluted using NGM buffer in a series of gradients (supernatant: NGM buffer = 1:1, 1:1.5, 1:2, 1:2,5, 1:3, 1:3.5) with a total volume of 1mL each. Diluted PA14 supernatants were exposed in the air for odorant evaporation for 5-6 hours before use. A drop of 10uL of diluted supernatant was placed in a fixed point on a 10-cm NGM assay plate immediately before a worm was placed 1.5 cm away from the drop. The tested worm was transferred to an empty NGM assay plate to remove the bacteria from its body before being transferred to the assay plate. The series of diluted PA14 supernatants was tested first to identify the most attractive dilution for the formal assay based on how efficiently naive worms navigated to the drop. Using the identified PA14 supernatant dilution, the formal experiment was recorded using a Grasshopper3-GS3-U3-120S6M camera at 7 fps. The recording stopped when the worm entered the supernatant drop or moved out of the recording view or at 5 min after recording started, whichever came first. The recorded chemotactic trajectory was analyzed using WormLab (MBF Bioscience) and a MATLAB (MathWorks) code to generate navigation index between the starting point and the drop position. The navigation index is defined as the ratio of radial speed (V_R_) and the locomotory speed (V_L_), and the navigation index of an assay is the mean value of the navigation indices calculated for every 2 s (14 frames). Statistical analysis was done using GraphPad Prism 10.

#### Slow-killing assay

Slow-killing assay was performed as previously described^84^ with minor modifications. OP50 and PA14 culture were prepared in NGM medium at 27 °C overnight. 0.5 mL of each culture was fully spread on each 6-cm NGM plates. The plates were incubated at 37 °C for 24 hours and then at 25 °C overnight before 20 young adult hermaphrodites were transferred to each plate. All assay plates were kept at 25 °C and scored for survived worms twice a day. Survived worms were identified when they responded to a gentle touch of a platinum picker. Survived worms were transferred to fresh slow killing assay plates every day. Five assay plates were scored per genotype per condition per assay. Pairwise Mantel–Cox log-rank test was used to compare each mutant strain with the reference wild-type strain, with replicate assay included as a stratification factor and Bonferroni multiple comparison test used for correction. All coding, statistical tests, and graphing were performed using R 4.5.1.

#### Calcium imaging

Calcium imaging and analysis were performed similarly as previously described^54,101^. PDMS microfluidic chips were used to restrain movements of the worms during time-lapse calcium imaging recording in worms. Individual naive or trained worms were transferred to an empty NGM plate to remove the bacteria from the body and then transferred to NGM buffer. A small syringe containing NGM buffer with 1% PLURONIC® F-68 (Gibco) was used to retrieve the worm from the NGM buffer and to load the worm into the microfluidic chip. Fresh OP50 (OD = 0.3 – 0.4) and PA14 (OD = 0.6 – 0.75) cultures were prepared by inoculating single bacteria colonies into NGM medium and incubating at 26-27 °C overnight. After centrifugation of the fresh cultures at 3,000rpm for 14 minutes, OP50 and PA14 supernatants were added to the large syringes connected to the microfluidic chips to provide stimuli into the PDMS chip. Time-lapse fluorescence imaging was performed using a confocal Nikon Eclipse Ti-E inverted microscope with a 40x oil-immersion objective, a Yokogawa CSU-X1 scanner unit, an ANDOR iXon Ultra EMCCD camera and NIS-Elements (version AR 4.13.04) at 5 fps. For all recordings, the stimuli were given in a standard temporal pattern: OP50 (60 seconds) – PA14 (30 seconds) – OP50 (30 seconds). In each worm, 1-5 focal planes containing neurons of interest were recorded separately for time-lapse GCaMP6 signals. For URY and OLQ neuron imaging, time-lapse nuclear-localized mCherry signals were also recorded to correct the effect of worm body movement on GCaMP6 signal intensity.

For analysis, because the first 30-second exposure to OP50 stimulation was used for worm to adapt to the imaging condition, the time-lapse image stacks of the remaining 90 seconds were aligned using the StackReg plugin (rigid body transformation) in ImageJ. Regions of interest (ROIs) of each neuron of interest were selected manually, and average GCaMP6 and NLS-mCherry-NLS signals were measured followed by the subtraction of background signals measured from a near region of the same size.

For calcium imaging data with only GCaMP6 signals, the average GCaMP6 signal for the first 30 seconds of OP50 stimulation was defined as baseline (F_b_) and the change of GCaMP6 signal relative to the baseline was calculated as follows:

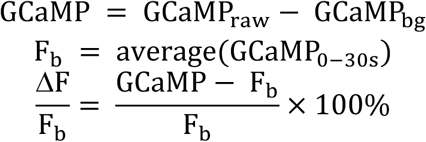

where GCaMP_raw_, GCaMP_bg_, are mean values of the raw GCaMP6 neuronal and background fluorescence intensity signals, respectively.

For calcium imaging data with both GCaMP6 and nuclear-localized mCherry signals, the ratiometric fluorescence intensity R was calculated and further normalized by the ratiometric fluorescence intensity R of the first 30s as follows:

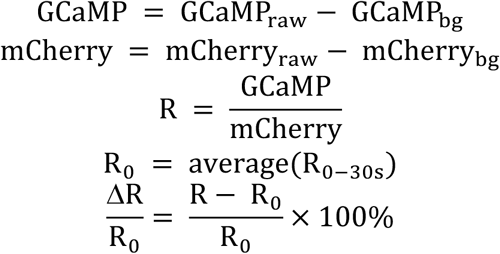

where GCaMP_raw_, mCherry_raw_, GCaMP_bg_, mCherry_bg_ are the mean values of raw GCaMP6 and mCherry neuronal and background fluorescence intensity signals, respectively.

Several tested neurons include left/right subtypes and dorsal/ventral subtypes, and recordings for left/right subtypes were merged for data analysis. Averaged 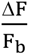 or 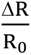 of each neuron during PA14 stimuli was calculated followed by statistical analysis to compare the difference between naive and trained conditions in each genotype. Statistical analysis was done using GraphPad Prism 10.

#### Confocal microscopy

Worms were anesthetized with 0.1 mM sodium azide and placed on a 2% agarose pad covered with 0.16-0.19 mm cover slip. Fluorescence signals from the worms were captured by performing Z-stack images using a confocal Nikon Eclipse Ti-E inverted microscope with a 40x oil-immersion objective or 4x air objective, an ANDOR iXon Ultra EMCCD camera and NIS-Elements (version AR 4.13.04). For learning-associated DAS gene expression, a maximum intensity projection of each channel for each image was generated. The brightness and contrast were adjusted for each channel in each image to aid the visualization of fluorescent signals. Montages containing the single channels and merged channel were generated using ImageJ for display.

To quantify UPR^MT^ and PA14-induced immune response, two naive or trained worms expressing the GFP reporters were aligned together for Z-stack image recording under a 4x air objective with 1.5x zoom-in of the field of view. To exclude the interference from worm autofluorescence, control groups where worms without the GFP reporters but in the same genetic backgrounds were also imaged in parallel. To quantify fluorescence signals, a maximum intensity projection of each image was generated, and average signal intensities of the regions of interest (ROIs) over the 2 aligned worms and the image background of the same area size were measured using ImageJ. The GFP and autofluorescence signals in each image of each condition for each strain were calculated as follows:

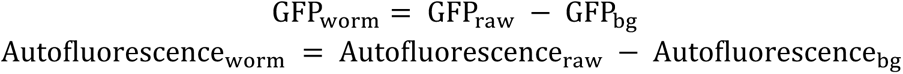

where GFP_raw_ and GFP_bg_ are the mean values of the raw GFP and background fluorescence in the GFP images, respectively; Autofluorescence_raw_ and Autofluorescence_bg_ are the mean values of the raw autofluorescence and background fluorescence in the control images, respectively.

The net GFP signals from the GFP reporters in each condition for each strain were calculated as the difference between the GFP signal of each recoding and averaged autofluorescence signals from the images taken in parallel on the same day:

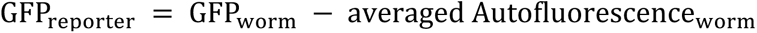

This was followed by wild-type GFP_reporter_ normalization by normalizing the GFP_reporter_ signals from individual images to averaged GFP_reporter_ signals from wild-type worms under the naïve condition for each replicate. Statistical analysis was done using GraphPad Prism 10.

## SUPPLEMENTARY INFORMATION

Document S1. Figures S1-S6

Table S1. Genes with differential transcript abundance (FDR < 0.05) for all pairwise comparisons, related to Figure 1

Table S2. LSVs with differential AS (|dPSI| > 0.2, probability of changing > 0.6) for all pairwise comparisons, related to Figure 1

Table S3. Learning-associated DT (differential transcript abundance) genes under threshold FDR < 0.05 and |log_2_FC| > 1, related to Figure 2

Table S4. Learning-associated DT (differential transcript abundance) genes under threshold FDR < 0.01 and |log_2_FC| > 3.5, related to Figures 2, S3 and S4

Table S5. Learning associated DAS (differential alternative splicing) LSVs under threshold |dPSI| > 0.2, probability of changing > 0.6, related to Figures 2, S3 and S4

Table S6. Transgenic and mutant worms, plasmids and primers used in this study

## ACKNOWLEDGEMENT

We thank *Caenorhabditis Genetics Center* (supported by the NIH Office of Research Infrastructure Programs P40 OD010440) and National BioResources Project (NBRP) (supported by Japanese government) for some strains used in this study; SunyBiotech for generating the CRISPR-engineered *twnk-1* alleles *syb9558* and *syb10064*; Drs. C. Bargmann, O. Hobert, E. Yemini for transgenes; X. Gracida and T. Wu for their guidance on applying TRAP-RNAseq technique; the Bauer Core Facility at Harvard University for providing the sequencing service; the FASRC Cannon cluster supported by the FAS Division of Science Research Computing Group at Harvard University for part of sequencing data analysis; and Zhang laboratory members for discussions on this project. This study is partly supported by the National Institute of Health (R21NS121825) and Harvard-/MIT Joint Research Grants Program in Basic Neuroscience.

## AUTHOR CONTRIBUTIONS

Project conception, experiment design and result interpretation: M.C., J.C., and Y.Z.

Sequencing data collection and analysis: M.C., M.W., and B.K.

Functional characterization data collection and analysis: M.C.

Transgene and transgenic strain constructions: M.C., J.L., and M.G.

M.C., J.C. and Y.Z. wrote the manuscript with input from other authors.

---

**Figure S1.**
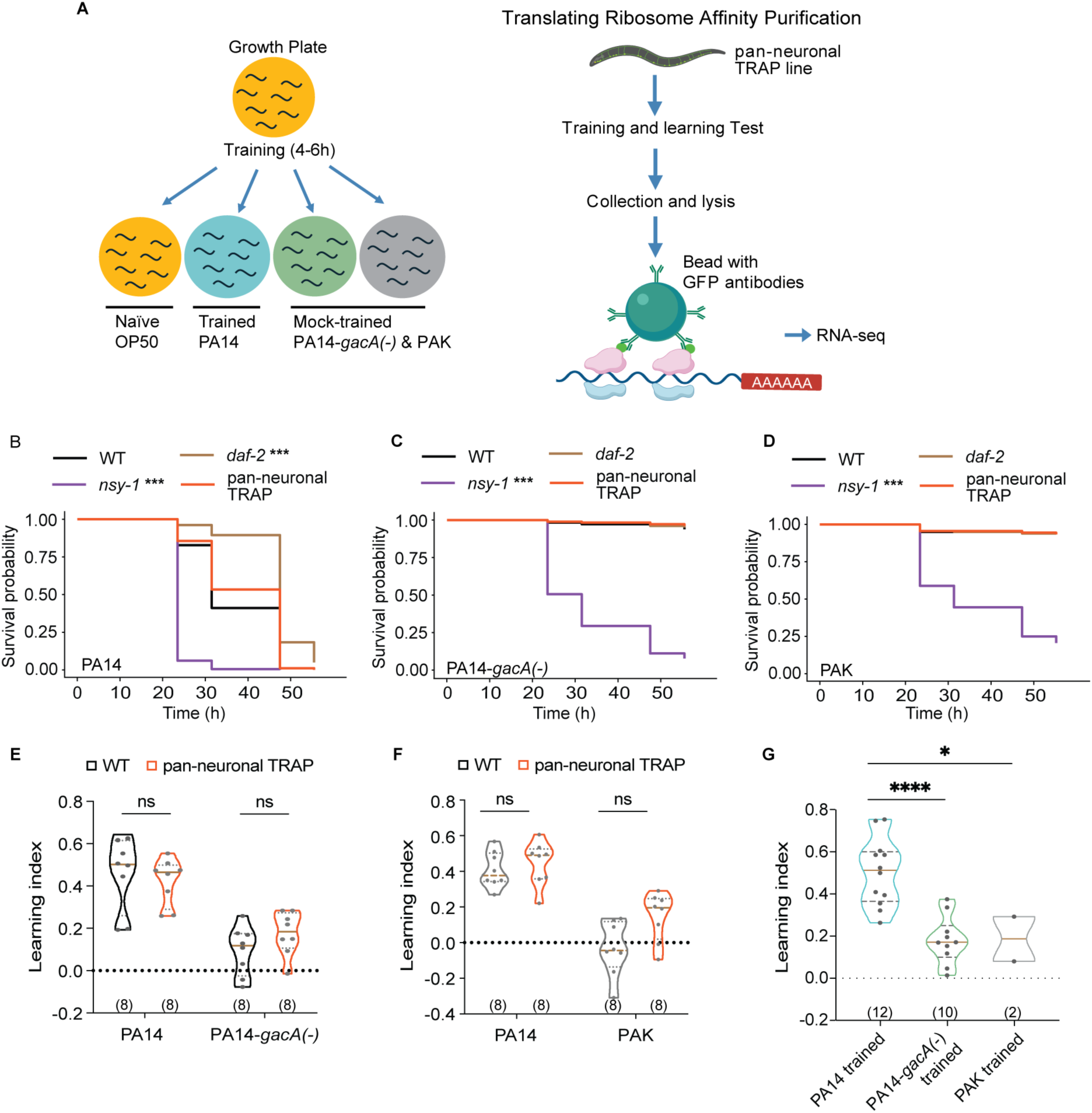
TRAP-RNAseq experiment, related to Figure 1. (A) Experimental outline for aversive training process and TRAP-RNAseq. Worms pan-neuronally expressing a GFP-tagged ribosomal protein in the nervous system were trained in four conditions for 4-6 hours. Worm lysates were immunoprecipitated with anti-GFP antibodies and neuronal ribosome-associated-mRNAs were collected for sequencing. Partly created in BioRender. Chen, M. (2026) https://BioRender.com/utzmvtn. (B-D) Kaplan–Meier survival curve of wild-type, pan-neuronal TRAP worms in a slow-killing assay of PA14, PA14-*gacA(-)* and PAK over two days. Pairwise Mantel–Cox log-rank tests compared each mutant strain with wild type, with experimental replicate included as a stratification factor and Bonferroni method as multiple comparison test. *** p < 0.001. Summarized data from 2 independent assays are shown. 100 or 80 animals were used per genotype per assay. (E-F) Violin plots of learning indices measured by droplet assay in wild-type worms and pan-neuronal TRAP worms in the PA14-trained and PA14-*gacA(-)*-trained (E), or PA14-trained and PAK-trained conditions (F). (G) Violin plots of learning indices measured by droplet assay in pan-neuronal TRAP worms in the PA14-trained, PA14-*gacA(-)*-trained and PAK-trained conditions during sample collections for TRAP-RNAseq experiment. For (E-G), violin plots demonstrate individual data points and frequency distribution with horizontal lines illustrating median (brown solid) and quartiles (gray dashed). Multiple unpaired t tests between wild-type and pan-neuronal TRAP worms across training bacteria, with Holm-Sidak multiple comparison test (E and F); or one-way ANOVA with Dunnett multiple comparison test (G). * p<0.05, **** p < 0.0001, ns, not significant. Numbers in parentheses indicate number of independent assays for each condition. Summarized data from 4 independent days (E-G).

**Figure S2.**
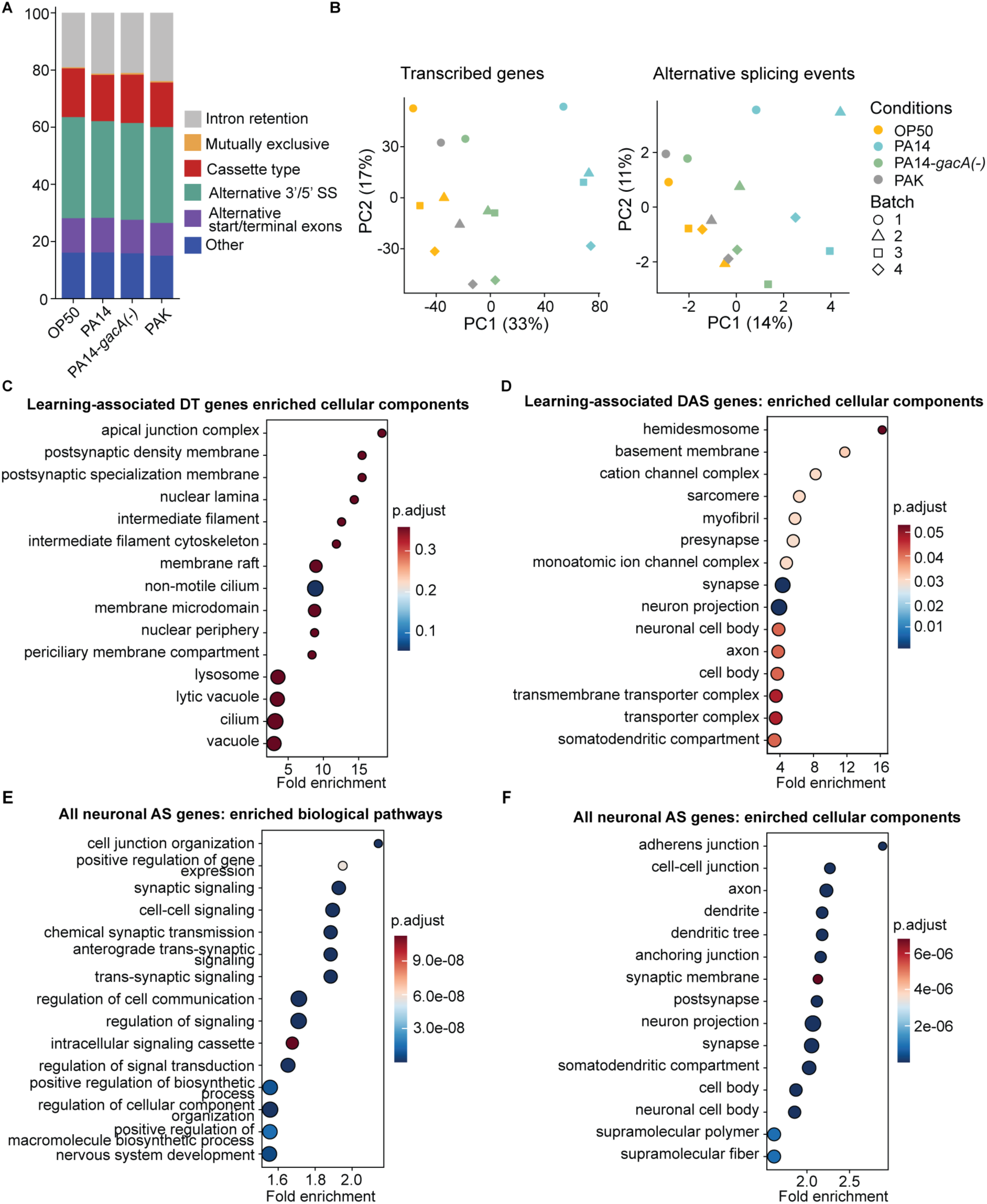
Aversive experience regulates both transcript abundance and alternative splicing (AS) globally, with each operating on functionally distinct gene sets, related to Figure 2. (A) Relative proportion of all neuronal AS LSVs from all four conditions classified into five canonical splicing types and an additional ambiguous type. (B) PCA analysis on all neuronally expressed genes (left) and all neuronal AS LSVs (right) across all samples with the outlier sample PAK3 removed, respectively. The batch effect was obvious in PCA plot using all neuronally expressed genes but not PCA plot using all neuronal AS LSVs. Gene expression or splicing levels were centered but not scaled. (C, D) Enriched cellular components of learning-associated DT and DAS genes relative to all neuronally expressed genes. (E, F) Enriched biological pathways and cellular components of all neuronal AS genes relative to all neuronally expressed genes. For (C-F), Gene Ontology (GO) analysis was performed applying Benjamini-Hochberg (BH) method for multiple comparisons. The enrichment ratio is the frequency of input genes to the background frequency of genes in each GO term. Significantly enriched GO terms with p.adjust < 0.1 and top 15 enrichment ratios are shown.

**Figure S3.**
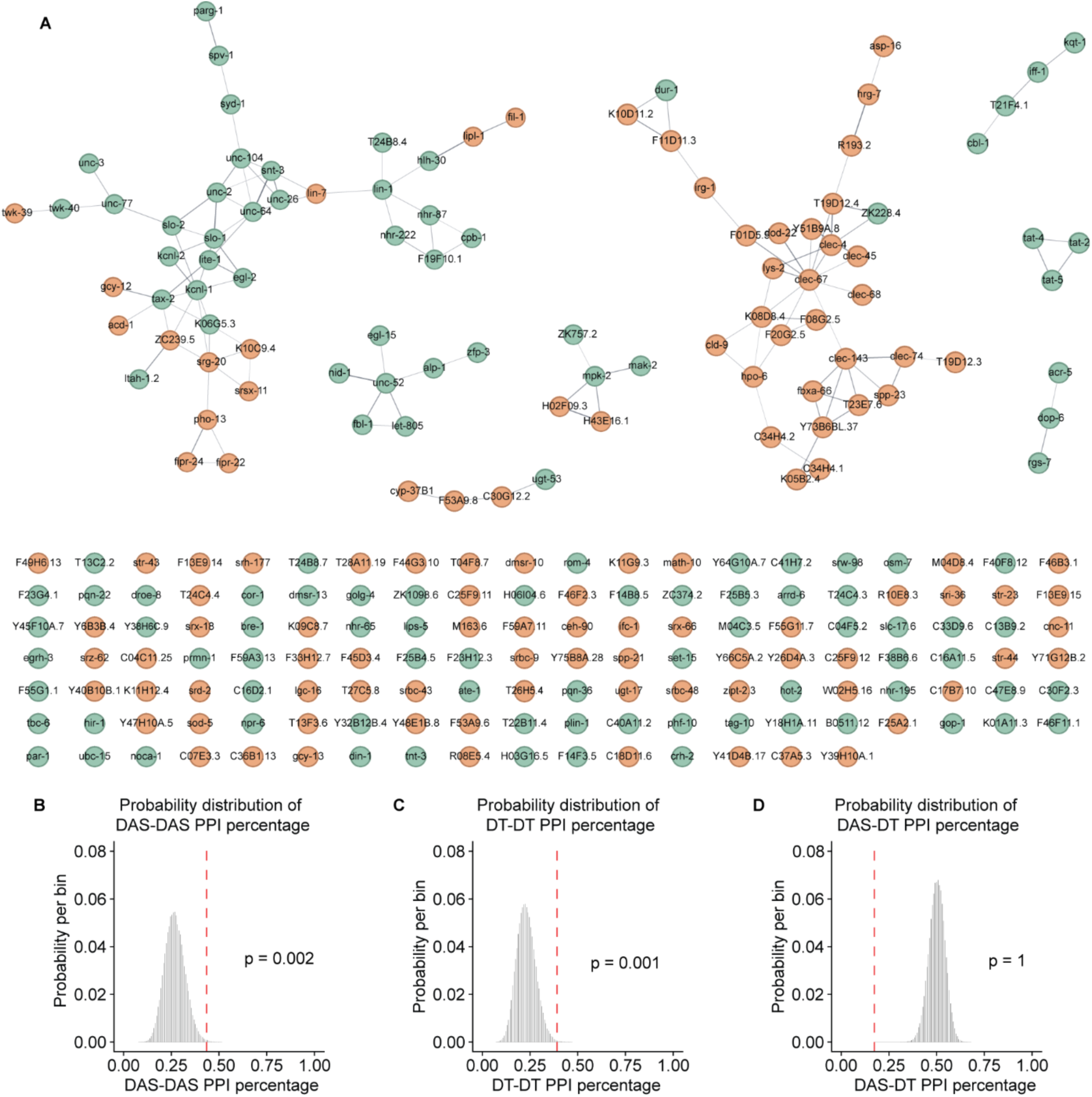
PPI (protein-protein interaction) network of learning-associated DT and AS genes shows the enriched interactions within each gene set, related to Figure 2. (A) PPI network of learning-associated DT and DAS genes from STRING database. Green circles denote genes only showing differential AS and orange circles demote genes only showing differential transcript abundance. (B-D) Probability distribution of PPI percentages for AS-AS, DT-DT and AS-DT interactions during 100,000 times of permutation. Red dashed lines denote the observed probability for each interaction type in the real network.

**Figure S4.**
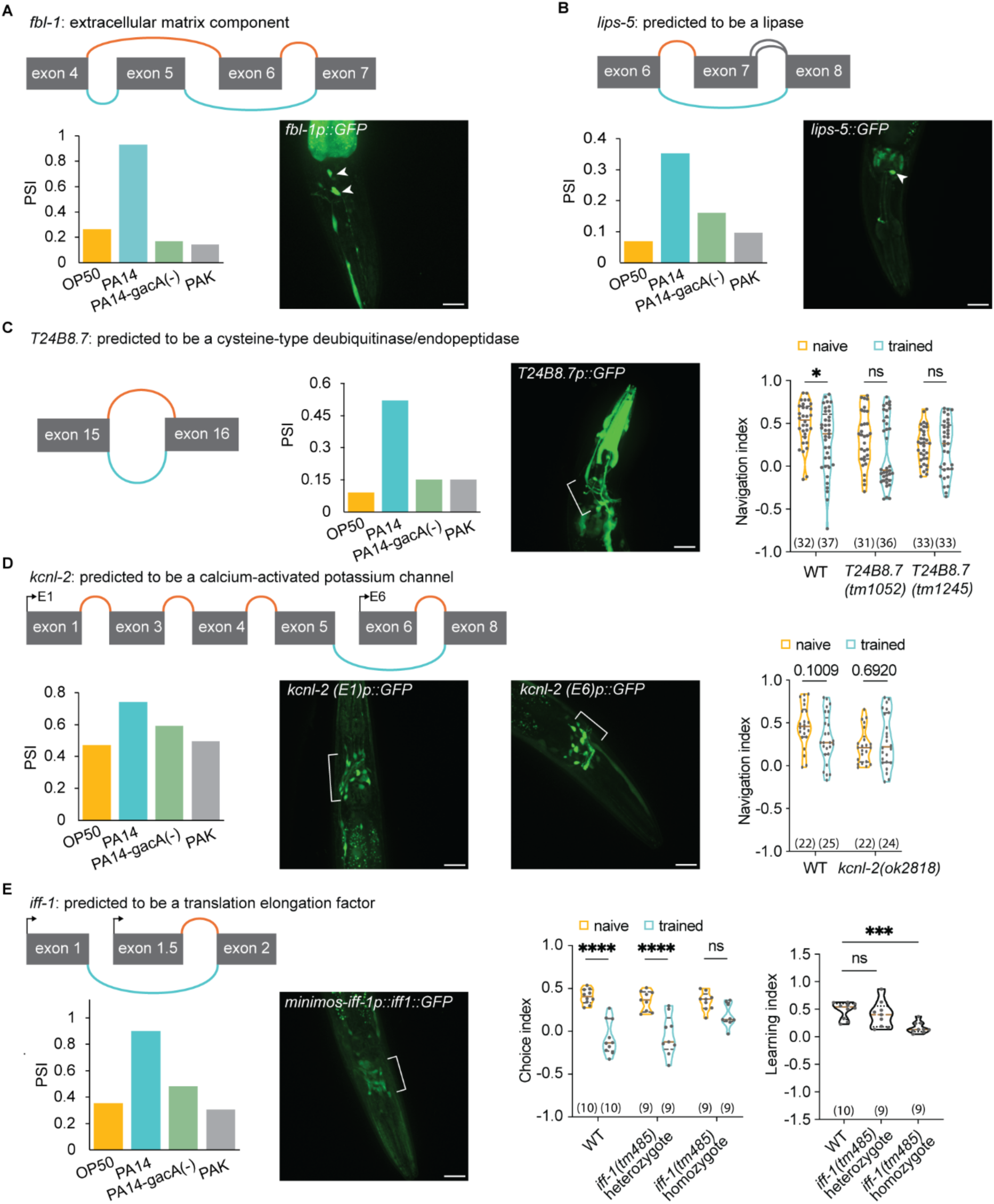
Top learning-associated DAS identify neuronally expressed genes that regulate aversive learning, related to Figure 3. (A, B) Learning-associated DAS events (up) in *fbl-1* (A) and *lips-5*(B), their splicing levels in all four conditions (bottom left) and reporter-based gene expression pattern (bottom right). Arrow heads indicate neurons expressing the candidate genes. Scale bars, 20 µm. (C, D) Learning-associated DAS events (up), splicing levels in all four conditions (bottom left), expression pattern (bottom center) and violin plots of navigation indices towards PA14 supernatant under naive and PA14-trained conditions in wild-type worms and in worms containing mutation in gene candidates (bottom right) for identified learning-associated DAS candidates *T24B8.7* (C) and *kcnl-2* (D). Brackets indicate the enriched neuronal expression of the candidate genes. (E) Learning-associated DAS events (up), splicing levels in all four conditions (bottom left), expression pattern (bottom center) and violin plots of choice indices and learning indices measured by droplet assay in wild-type worms and the mutant worms (bottom right) for the identified learning-associated DAS candidate *iff-1*. For (A-E), for each gene only the genomic regions involved in the learning-associated AS events are shown. Orange representing the junctions downregulated, blue representing the junctions upregulated, and gray representing the junctions not significantly changed after PA14 training. For (C-E), violin plots demonstrate individual data points and frequency distribution with horizontal lines illustrating median (brown solid) and quartiles (gray dashed). Two-way ANOVA was used to compare different conditions within each genotype, with Sidak multiple comparison test (C, D and Choice index in E). One-way ANOVA was used to compare learning indices of mutant worms with wild-type worms, with Dunnett multiple comparison test (Learning index in E). * p<0.05, *** p < 0.001, **** p<0.0001, ns, not significant. Numbers in parentheses indicate number of worms (C, D) or number of independent assays (E). Data was collected from 4 (C) or 3 (D) independent assays or from 3 different days (E). White brackets indicate regions of neurons expressing the candidate genes. Scale bars, 20 µm.

**Figure S5.**
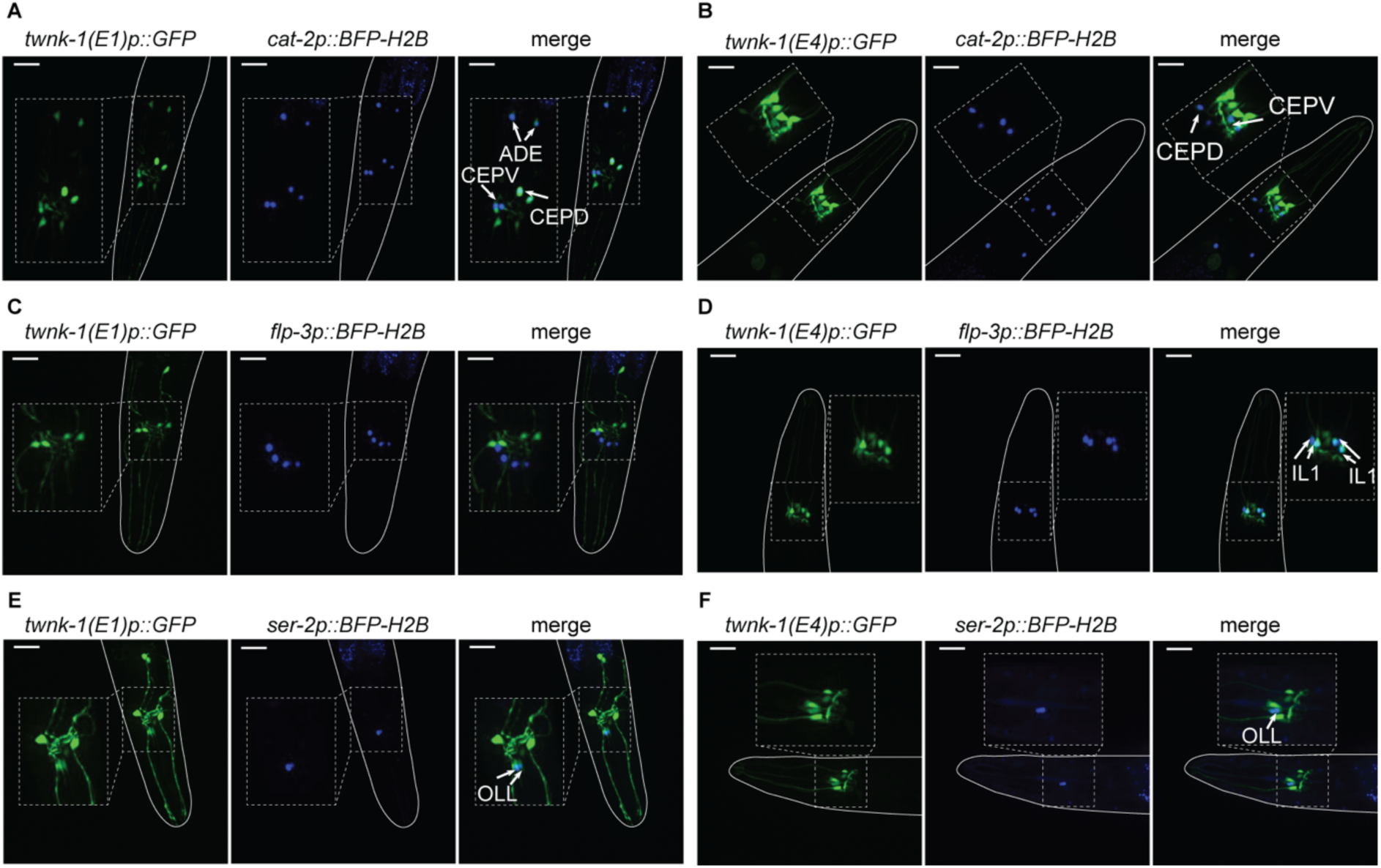
Colocalization of neuron-specific nuclear-localized BFP reporters with *twnk-1* GFP reporter identified neurons expressing *twnk-1* isoforms, related to Figure 3 and also see Figures S7C-S7H. (A-F) Representative images of co-localization of *cat-2p::BFP-H2B*, *flp-3p::BFP-H2B*, *ser-2p::BFP-H2B* with *twnk-1(E1)p::GFP* or *twnk-1(E4)p::GFP* reporters in naive adult worms.. Dashed lines outline the enlarged regions of interest. Lines outline the imaged worms. Arrows indicate the neurons co-expressing BFP and GFP with the neuron IDs specified. Scale bars, 20 µm.

**Figure S6.**
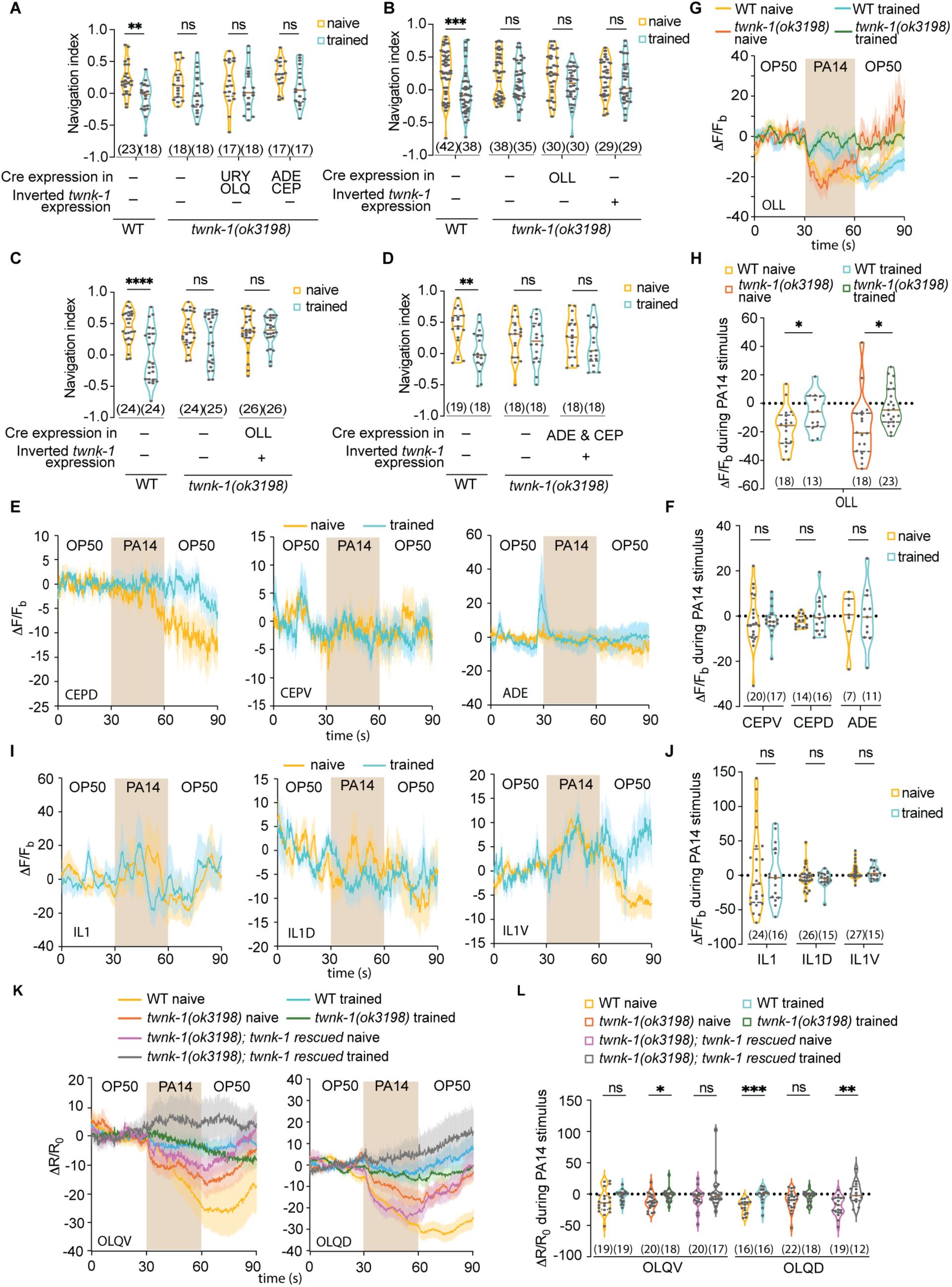
*twnk-1* expression in ADE, CEP, OLL or IL1 is not sufficient for aversive learning, related to Figure 4. (A-D) Violin plots of navigation indices towards PA14 supernatant under naive and PA14-trained conditions in wild-type worms, *twnk-1(ok3198)* mutant strain, and control transgenic *twnk-1(ok3198)* mutant worms carrying inverted *twnk-1* or *cre* transgene only (A, B), or in wild-type worms, *twnk-1(ok3198)* mutant strain, and transgenic *twnk-1(ok3198)* mutant worms expressing *twnk-1* DNA selectively in ADE and CEP neurons (C) or in OLL (D). (E) Traces of GCaMP6 signals in CEPV (left), CEPD (middle), and ADE (right) neurons in response to OP50-PA14-OP50 odorant stimulation in wild-type worms under naive and PA14-trained conditions. (G) Traces of GCaMP6 signals in OLL neurons in response to OP50-PA14-OP50 odorant stimulation in wild-type worms, and *twnk-1(ok3198)* mutant worms under naive and PA14-trained conditions. (I) Traces of GCaMP6 signals in IL1 (left), IL1D (middle), and IL1V (right) neurons in response to OP50-PA14-OP50 odorant stimulation in wild-type worms under naive and PA14-trained conditions. (K) Traces of GCaMP6 signals in OLQV (left) and OLQD (right) neurons in response to OP50-PA14-OP50 odor stimuli in wild-type worms, *twnk-1(ok3198)* mutant strains, and transgenic *twnk-1(ok3198)* mutant worms carrying the full-length genomic *twnk-1* DNA under naive and PA14-trained conditions. For (E, G, I, K), lines in traces, mean; shades in traces, s.e.m.; brown rectangular shade, PA14 stimulus. ι1F/F_b_ and ι1R/R_0_, see Methods. For (F, H, J, L), violin plots of quantification of average GCaMP6 signals of 30s PA14 stimulus as shown in (E), (G), (I), and (K), respectively. For (A-D, F, H, J, L), violin plots demonstrate individual data points and frequency distribution with horizontal lines illustrating median (brown solid) and quartiles (gray dashed). Two-way ANOVA was used to compare conditions within each genotype, with Sidak multiple comparison test (A-D); or multiple unpaired t-tests were used to compare naive and trained conditions for all genotypes, with Holm-Sidak multiple comparison test (F, H); or multiple Mann-Whitney tests (J, L) between naive and trained conditions across genotypes, with Holm-Sidak multiple comparison test. * p<0.05, ** p < 0.01, *** p < 0.001, ns, not significant. Numbers in parentheses indicate number of worms assayed (A-D) or number of each neuron type recorded in each condition (F, H, J, L). Data was collected from 3 (A, C, F, H, J) or 6 (B) or 4 (D, I) independent assays.

**Figure S7.**
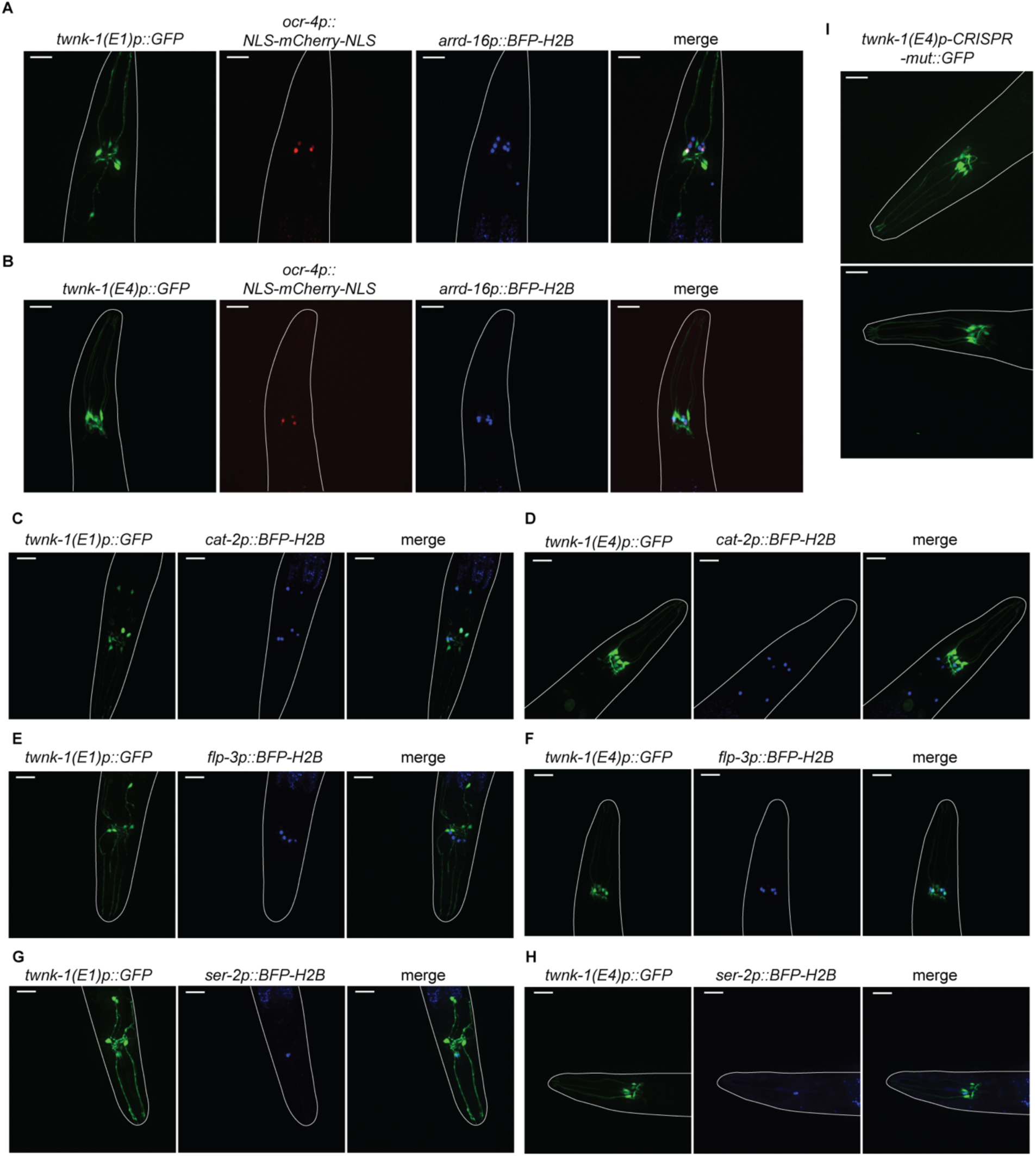
Microscopy images of expression patterns of GFP reporters driven by *twnk-1* promoters, related to Figures 3E-F and S5. (A, B) Raw images of Figures 3E and 3F. (C-H) Raw images of Figure S5. (I) Representative images of the expression pattern of a GFP reporter driven by the promoter of *twnk-1* E4 isoforms containing the same CRISPR-engineered mutations from *twnk-1(syb10064)*. Lines outline the imaged worms. Scale bars, 20 µm.

